# Dextran-based T-cell expansion nanoparticles for manufacturing CAR T cells with augmented efficacy

**DOI:** 10.1101/2025.04.10.648181

**Authors:** Tao Zheng, Keerthana Ramanathan, Maria Ormhøj, Mikkel Rasmus Hansen, Hólmfridur Rósa Halldórsdóttir, Hanxi Li, Kamilla Kjærgaard Munk, Carlos Rodriguez-Pardo, Rasmus Ulslev Wegener Friis, Seder Fayyad Islam, Peter M. H. Heegaard, Klaus Qvortrup, Hinrich Abken, Yi Sun, Sine Reker Hardup

## Abstract

Adoptive T cell therapy (ACT) using chimeric antigen receptor (CAR) engineered T cells is currently being explored in multiple cancer types beyond leukemia/lymphoma. A key step in CAR-T cell manufacturing is the activation and expansion of T cells, which facilitates viral transduction, however, may hamper T cell fitness and reduce *in vivo* persistence. We developed “T-Expand” for T cell activation and expansion, comprising dextran-based nanoparticles (NPs) conjugated with anti-CD3 and anti-CD28 antibodies. The NPs triggered robust polyclonal expansion of human T cells with efficiency in the range of commercial microbeads (Dynabeads™). Engineered in presence of T-Expand, CD19 CAR T cells exhibited enhanced proliferative capacity, cytotoxicity and persistence in *vitro*, and furthermore, showed superior anti-lymphoma activity in mouse models resulting in complete tumor clearance at one fourth of the CAR T cell dose. Importantly, T-Expand is biocompatible with no observed toxicity, circumventing removal steps after T cell expansion compared to Dynabeads^TM^. As a biocompatible T cell expansion platform, T-Expand simplifies the manufacturing process while enhancing T cell persistence and functionality, thereby holding promise for increasing clinical efficacy of CAR T cell therapy.

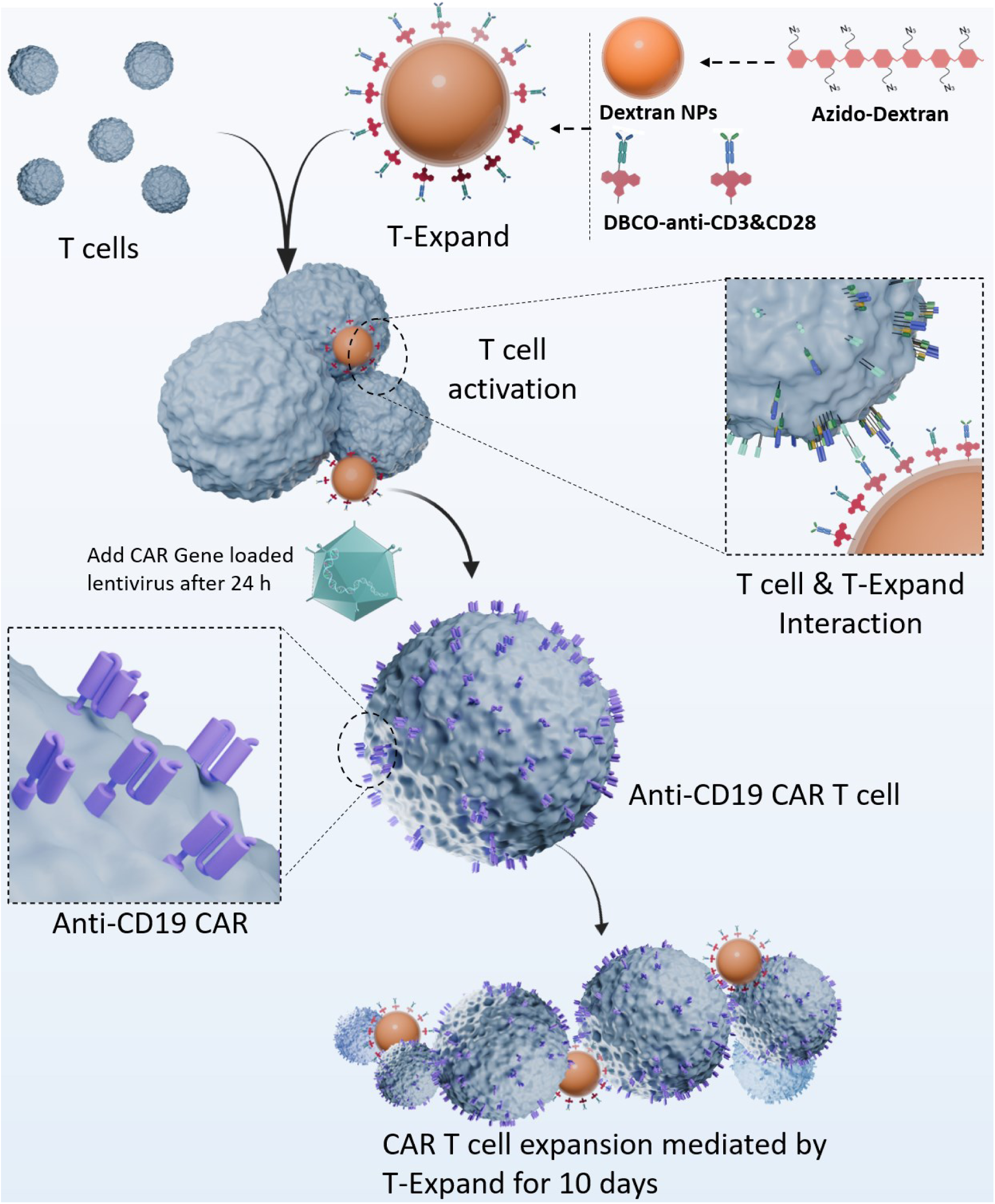
Graphical abstract/Cover figure: Illustration of CAR T cell manufacturing using T-Expands *ex vivo*.

## Introduction

T cells genetically modified to express a chimeric antigen receptor (CAR) has shown tremendous potential against a variety of hematological malignancies(*1, 2*). Despite the clinical success, a substantial number of patients do not gain long-term treatment benefit, potentially owing to T cell intrinsic factors leading to lack of functional capacity and in vivo persistence(*3, 4*). While the genetic modification of T cells requires potent T cell activation to enter a proliferative state susceptible to gene modification, the expansion phase imprints the functional characteristics of the resulting CAR T cells(*5*).

Current conventional CAR T-cell expansion platforms are based on immobilized anti-CD3 and anti-CD28 antibodies on magnetic microbeads (Dynabeads^TM^) or on a polymer matrix (TransAct^TM^)(*6, 7*). However, these artificial antigen-presenting cell (aAPC) platforms can lead to overstimulation of T cells causing cell exhaustion and ultimately result in lack of persistence after infusion(*8*). Alternative microscale aAPC platforms such as lipid-coated mesoporous silica scaffolds promote T cell expansion and demonstrated anti-tumor activity in vivo(*9, 10*). Recently, aAPC platforms based on graphene oxide flakes and viscoelastic microgel have been reported(*11, 12*). These platforms could enhance expansion kinetics as compared to conventional platforms. However, graphene oxide based aAPCs possess a technical challenge when isolating expanded T cell products from scaffolds, which may prevent generation of therapeutic CAR T cell products(*9*). Furthermore, the microgel-based aAPC platforms have difficulty reproducing and maintaining mechanical properties when used at large scale, thus posing challenges for scale-up manufacturing(*13*).

To address these limitations, we designed a novel dextran-based nano-aAPC, called T-Expand, comprising dextran NPs decorated with anti-CD3 and anti-CD28 antibody on its surface. Dextran presents notable advantages, such as high density of modifiable functional groups, which enables conjugation of high amount of T cell stimulatory molecules. We hypothesize that enhancing the surface density of these stimulatory molecules could effectively compensate for the reduced physical contact area inherent to nanoscale aAPCs, thereby maintaining robust T-cell expansion. Meanwhile, we hypothesize that the avidity between T-Expand and T cells is not as strong as that between microbeads and T cells. Upon T-cell activation, T-Expand is either internalized into T cells with TCR or detached from the T-cell surface, which may help prevent T-cell exhaustion by avoiding prolonged stimulation. In addition, T-Expand is biocompatible and mechanically stable, thereby avoiding the extra separation step and is suitable for large-scale production.

We demonstrate that T-Expand offers an alternative approach to enhance the quality of CAR T-cell products and simplify the CAR T-cell manufacturing process with beneficial functionality. CAR T cells expanded with T-Expand exhibit significantly improved persistence and therapeutic efficacy in both *in vitro* and *in vivo* settings. Specifically, CD19 CAR T cells expanded using T-Expand display reduced exhaustion markers compared to those generated using Dynabeads^TM^. Additionally, T-Expand-produced CAR T cells are applicable for long-term tumor clearance, underscoring their potential clinical advantages over conventional aAPC platforms.

## Results

### Characterization of the T-Expand

Dextran was selected as the material for T-Expand due to its biodegradability, biocompatibility, abundant functional groups and ease of chemical modification(*14–16*). Herein, we modified the dextran polymer with multiple copies of azido groups, which enabled flexible conjugation of antibodies in the aqueous phase via bio-orthogonal approaches such as azido-alkyne cycloaddition and Staudinger ligation (**Figure 1a**). Firstly, the azido linker was activated using carbonyldiimidazole (CDI) (**Supplementary Fig. 1-Fig. 4**). Subsequently, the CDI-functionalized azido linker was reacted with the dextran to produce azido-modified dextran polymer (**Supplementary Fig. 5 and Fig. 6**), which was then acetylated to form the acid-sensitive azido-dextran polymer (**Supplementary Fig. 7**). The resulting hydrophobic azido-dextran NPs (“naked” NPs) were prepared by double emulsion technology. Measurement by NP tracking analysis (NTA) system showed that the naked NPs had a homogenous size distribution of about 150 nm (**Supplementary Fig. 8a, b**).

**Fig. 1.**
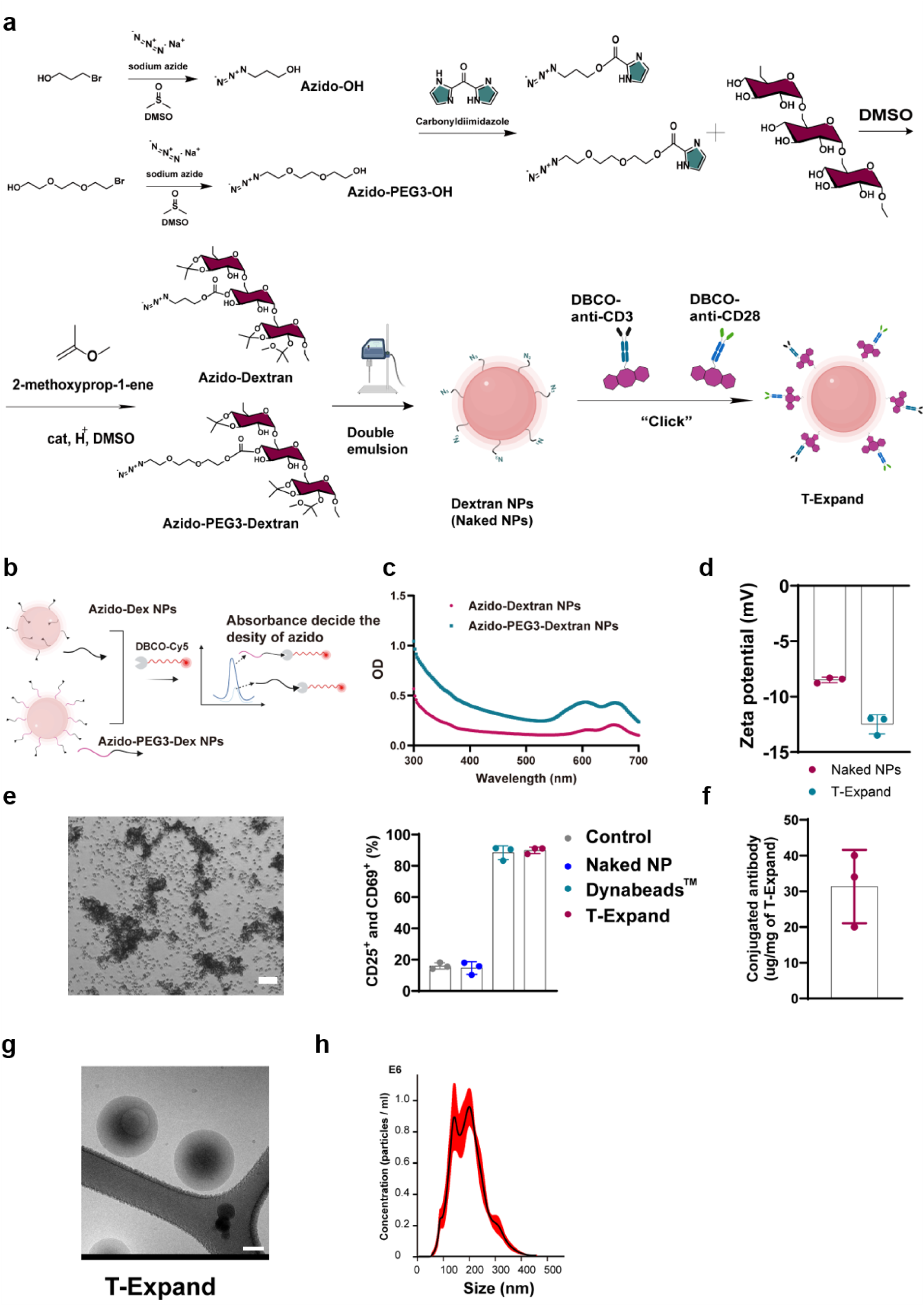
Design and characterization of anti-CD3 and anti-CD28 conjugated dextran-based T-Expand. **a,** Synthesis of azido modified dextran polymer, and the fabrication of T-Expand. **b,** UV-spectrum confirming that the azido linker is primarily orientated on the surface of naked NPs. **c,** UV-spectrum showing azido decorated dextran NPs in solution clicked with DBCO-Cy5 to determine the density of azido linkers. **d,** Zeta potential of naked NPs and T-Expand (n=3). **e,** Representative brightfield microscopy images of primary T cells cultured with T-Expand, scale bar: 100 nm. Percent of T cells expressing the activation markers CD25 and CD69 after co-incubation with T-Expand (number of donors: n = 3). **f,** The antibody conjugation efficiency on the surface of naked NPs measured by Nanodrop. **g,** Cryogenic transmission electron microscopy visualizing of T-Expand, scale bar: 50 nm. **h,** Size measurement of T-Expand by nanoparticle tracking analysis (NTA).

The hydrophobicity of the azido linker plays an important role in determining the orientation of the azido groups during the water/oil/water emulsion formation(*17*). To investigate the effect of the linker on the amount of azido groups on the surface, we selected two types of linkers, hydrophobic azido linker and amphiphilic azido linker with three PEG segments, to modify the dextran backbone before synthesizing the NPs. The distribution and orientation of the two linkers were investigated using a click-based colorimetric labeling method. Specifically, Cy5 molecules with dibenzocyclooctyne (DBCO) groups were clicked onto the naked NPs via click chemistry, and the overall density of azido group was determined by measuring the optical density (OD) value of Cy5 on the NPs. Higher OD indicates a higher number of azido functional groups on the surface of naked NPs, whereas lower absorption suggests that more azido groups are oriented inside the NPs (**Figure 1b**). The amphipathic linker with PEG segments orients more at the water–organic solvent interface, producing a higher density of azido groups on the NP surface compared to NPs without PEG segments (**Figure 1c**). This linker was applied for further use. The available azido groups on the surface of NPs for antibody conjugation was 24.6 nmol per mg dextran NP as determined by UV-Vis spectroscopy analysis (**Supplementary Fig. 8c, d**).

Next, DBCO modified anti-CD3 and anti-CD28 antibodies were confirmed by nanodrop measurement, which showed the distinct absorption peak of DBCO from 300-320 nm, indicating that the DBCO linker has been conjugated to the antibody (**Supplementary Fig. 9a, b**). Furthermore, the DBCO modified antibodies were conjugated to the surface of the naked NPs through click chemistry. After the conjugation, the zeta potential decreased from −8.5 ± 0.25 to −12.5 ± 0.87 mV (**Figure 1d**).

To evaluate whether the conjugated anti-CD3 and anti-CD28 antibodies were successfully linked and retained their function for T cell activation, the T-Expand was co-cultured with primary T cells and analyzed the expression of activation markers (CD25 and CD69) via flow cytometry. T cells were activated following formation of T cell clusters visible by microscopy, with subsequent expression percentage of CD25 and CD69 similar to those induced by Dynabeads™ (**Figure 1e**). In contrast, the naked NPs did not activate T cells, demonstrating the role of the conjugated antibodies on T-Expand. Nanodrop quantification of antibody conjugation efficiency revealed that ∼31.3 ug of antibodies were conjugated per milligram of NPs (**Figure 1f, Supplementary Fig. 10**). Compared to current T cell expansion scaffolds, our T-Expand achieves the highest antibody coupling among reported nanoscale T cell expanders, even surpassing some micron-scale scaffolds **(Supplementary Table 1).** Additionally, cryogenic transmission electron microscopy visualization revealed that T-Expand exhibited high uniformity (**Figure 1g**) at size of 199.8 ± 70.7 nm (**Figure 1h**). Overall, these data demonstrated that the click chemistry-mediated antibody conjugation method effectively anchors antibodies onto the naked NPs.

Next, we explored the presence of serum on T-Expand performance since several T-cell expansion protocols requires serum supplementation. the proteomics analysis revealed that the click chemistry-mediated antibody conjugation method also limits the protein corona formation on the surface of T-Expand (**Supplementary Fig. 11, 12**) and show no functional compromise for T cell activation (**Supplementary Fig. 12h, i)**.

### Investigation of the interaction of T-Expand with T cells

To study the interaction T-Expand with T cells, we engineered Jurkat T cells to express a cathepsin L-mCherry fluorescent reporter. Cathepsin L is a lysosomal protein co-localizing with perforin and Granzyme B in T cell granules(*18*). Using the Jurkat cathepsin L-mCherry reporter system, we investigated the ability of T-Expand to activate T cells by viewing the movement of cathepsin L-mCherry granules. Confocal imaging of engineered Jurkat cells in 3D and simulation shows the mCherry-Cathepsin L reporter has been successfully expressed in Jurkat cells (**Figure 2a**). Colocalization imaging indicates that T-Expand effectively activates Jurkat cells, accompanied by the translocation of mCherry-Cathepsin L to the T-Expand contact site, ultimately resulting in colocalization. In contrast, the naked NPs do not directly interact with the Jurkat cells (**Figure 2b, c**). Furthermore, with the combination of a 3D perspective simulation in measuring the distance between the T-Expand and the granules suggests colocalization, underscoring the specificity of T-Expand in activating Jurkat cells (**Figure 2d**). The real time-3D imaging shows that the T-Expand forms a multicentric granules cluster that is different from the monocentric granule formed with Dynabeads™ (**Figure 2e, Supplementary Video 1&2**), indicating that T-Expand could activate Jurkat cells at multiple interfaces between T-Expand and Jurkat cells.

**Figure 2.**
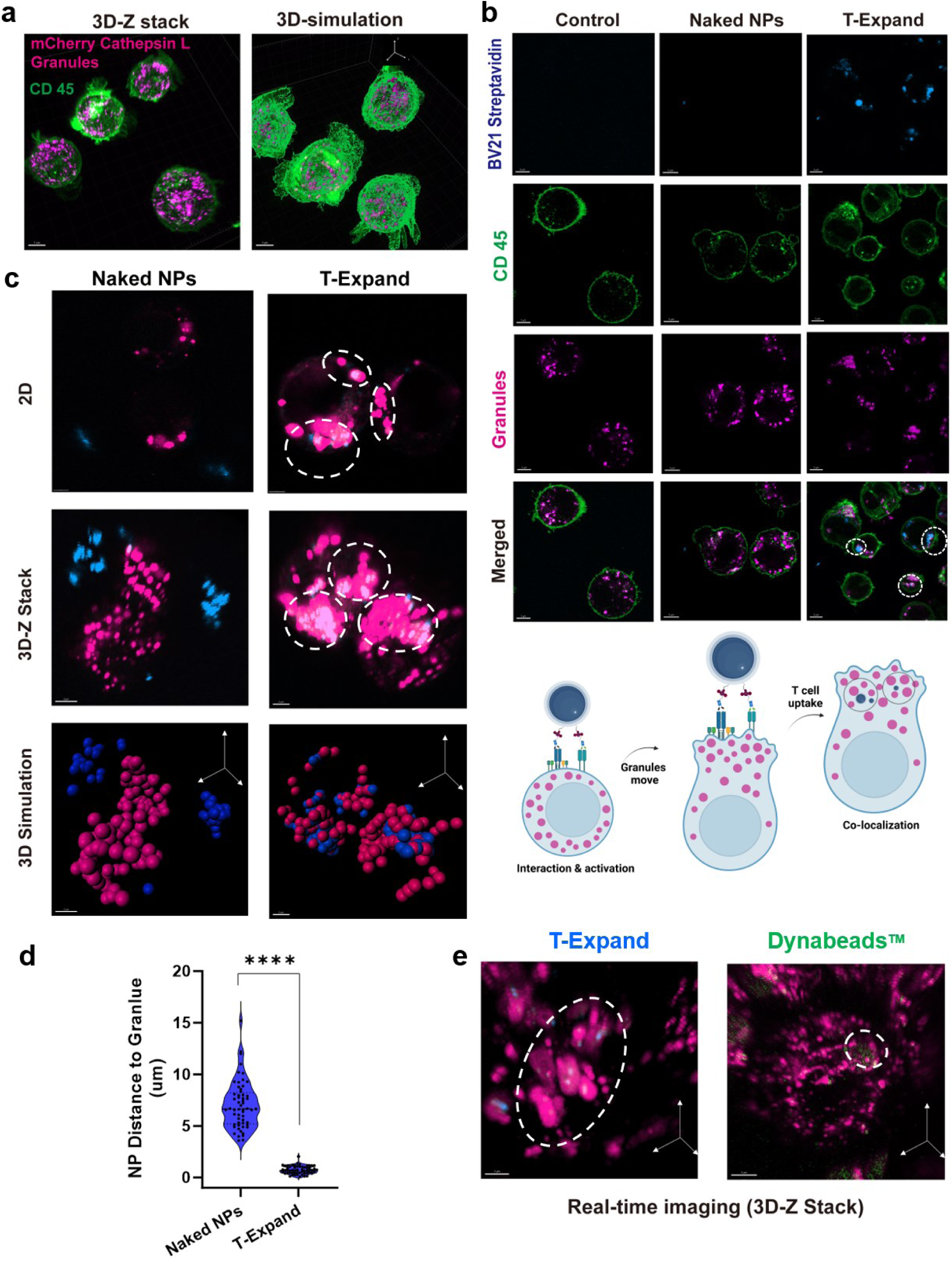
The investigation of T-Expand interacting with Jurkat cells. **a,** mCherry labelled Cathepsin L is located in Jurkat cell granules as shown by confocal imaging, scale bar: 5 um. **b,** Engineered Jurkats were incubated with T-Expand or naked NPs both loaded with BV421-streptavidin for 24 h and then imaged by Airyscan confocal imaging (2D-view), scale bar: 5 um. White circle indicates the location of naked NPs or T-Expand. **c,** Specific activation test, Airyscan imaging (3D view) and Imaris 3D simulation on how T-Expand interacting with engineered Jurkats, scale bar 2 um. **d,** The distance of naked NPs or T-Expand to mCherry-Cathepsin L under Imaris 3D simulation, (number of granules n = 130), ****P < 0.0001, two tailed unpaired t test. **e,** The real-time 3D confocal imaging indicates the granules cluster formation between Dynabeads™ and T-Expand. White circle indicates the location of T-Expand or Dynabeads^TM^, scale bar 3 um.

### Manufacturing of CAR T cells with T-Expand results in cell products with high proliferative capacity, cytotoxicity and persistence

We investigated two parameters known to influence T cell expansion; the ratio of conjugated anti-CD3: anti-CD28 antibodies on T-Expand and the dose of T-Expand. First, we investigated how the stoichiometric ratio of anti-CD3: anti-CD28 influences T cell activation. To avoid potential interference caused by variations in the concentration of anti-CD3 and anti-CD28, we ensured that the surface density of these factors was saturated in all experiments. Based on the T cell activation assay, the different stoichiometric ratios of anti-CD3: anti-CD28 antibodies did not influence significantly on the percentage cells positive for the activation markers CD25 and CD69. For all stoichiometric ratios, the percentage of these activation markers increased with higher concentrations of T-Expand, reaching saturation at 3.75 μg of T-Expand (0.75 mg/mL) per 0.5 × 10^6^ T cells (**Figure 3a**). At lower concentrations of T-Expand, we observed that T-Expand (anti-CD3: anti-CD28 ratio 1:3) demonstrated the highest fold expansion by day 14 **(Supplementary Figure 13a, b, c**).

**Figure 3.**
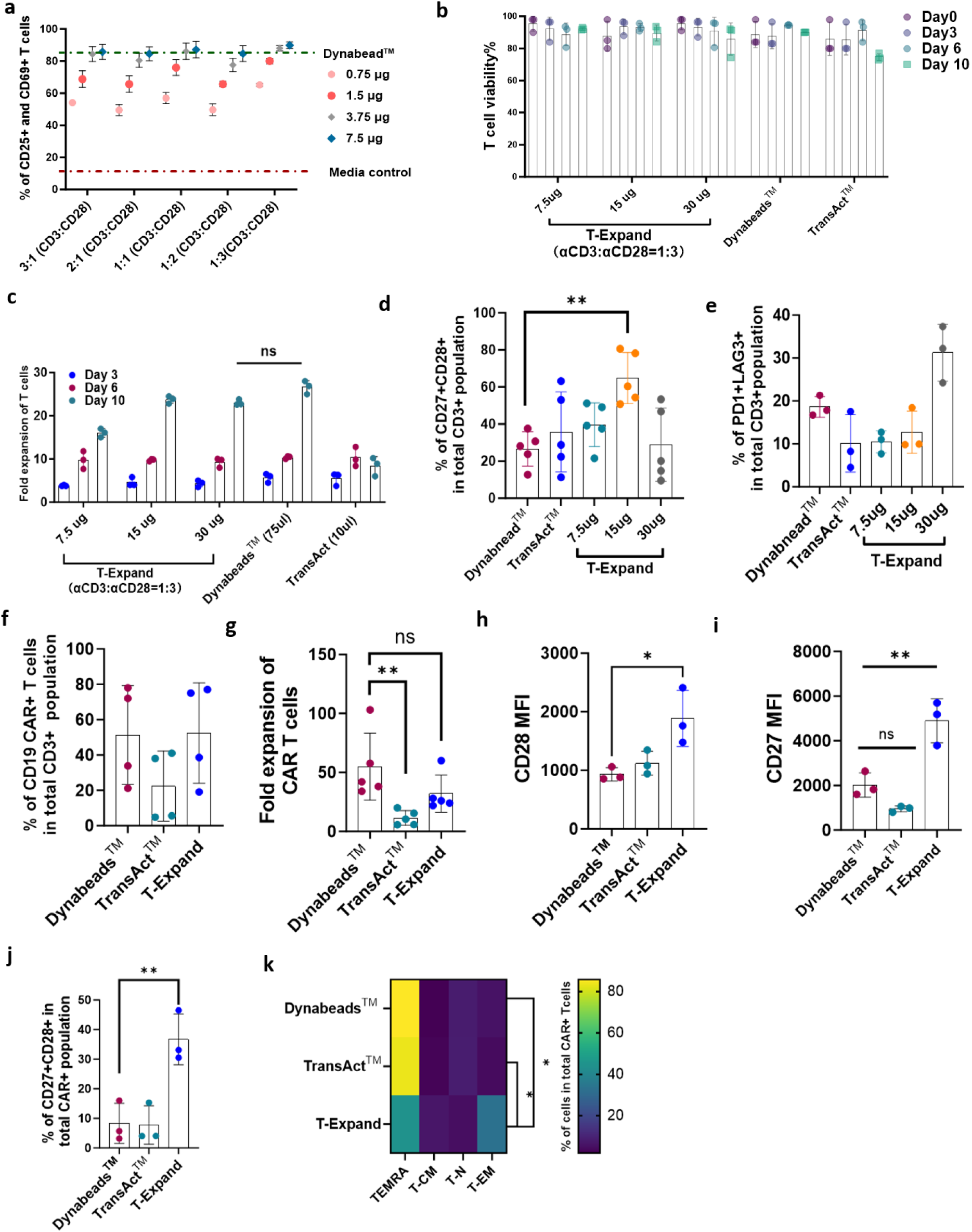
CAR T cell manufacturing with T-Expand results in T cell products with a favorable phenotype for ACT. **a,** T cell activation marker (CD25 and CD69) expression percentage as determined by flow cytometry following incubated with T-Expand with different ratio of anti-CD3: anti-CD28 for 24 h (number of donors: n = 3). **b,** A graph showing the viability of T cells on the different time points of expansion with different dose of T-Expand (or) Dynabeads^TM^ (or) TransAct^TM^; **c,** Graph showing the fold expansion of isolated T cells from peripheral blood expanded using T-Expand, Dynabeads™ or TransAct™. **d,** The expression CD27+ and CD28 on T cells expanded by T-Expand (conjugated anti-CD3: anti-CD28= 1:3) at different dose (7.5 ug, 15ug, 30 ug per million Pan T cells), Dynabead^TM^, and TransAct^TM^. **e,** The expression of the exhaustion markers of PD1+LAG3+co-expression on T cells expanded by T-Expand (anti-CD3: anti-CD28= 1:3) at different dose (7.5 ug, 15ug, 30 ug), Dynabeads^TM^, and TransAct^TM^. **f,** Percentage of CD19 CAR T cells in cultures after after lentiviral transduction and 10 days of expansion. **g,** Fold expansion of CAR T cells 10 days expansion with T-Expand, Dynabeads™ or TransAct™, Bargraph showing mean fluorescent intensity (MFI) of **h,** BUV737-CD28, and **i,** BV605-CD27on CAR T cells after expansion. **j,** Percentage of CD27+CD28+ CAR T cells after 10 days expansion with T-Expand, Dynabeads™ or TransAact™. **i,** Heat map of phenotype analysis (naïve T cell: TN; TCM, TEM, TEMRA) of expanded CAR T cell during10 days of expansion with T-Expand, Dynabeads™ or TransAct™. *Data in panels **d**, **e**, **f**, **g**, **h**, **i**, **j** and **k** represented as mean ± SD, of biological replicates with each dot indicating one healthy donor sample. Statistical significance determined via ordinary one-way ANOVA using Holm-Šídák’s multiple comparisons test *P < 0.05, **P < 0.01, ***P < 0.001, ****P < 0.0001. Data pannel in **c**, presented as mean ± SD, of biological replicates with each dot indicating one healthy donor sample. Statistical significance determined via two-way ANOVA using Dunnett’s multiple comparisons test*P < 0.05, **P < 0.01, ***P < 0.001, ****P < 0.0001*

Then we followed the viability and fold expansion of the T cells over a 10 day cultivation period and compared T-Expand with gold standard of Dynabeads™ and TransAct™ as the standards. (**Figure 3b-c**). T-Expand at the dose of 15 µg induced T cell expansion at a similar degree as Dynabeads™ by day 10 and superior to TransAct™ (**Figure 3c**). The T-Expand-expanded T cells predominantly consisted of 35-50% effector memory T cells (TEM) and approximately 40% effector memory T cells re-expressing CD45RA (TEMRA), which mirrored the distribution observed in Dynabeads^TM^-expanded T cells (**Supplementary Figure 13d**). Notably, this condition yielded higher CD27+CD28+ expression (**Figure 3d**) and lower levels of PD1+ and LAG3+ on T cells (**Figure 3e**). Consequently, the dose at 15 ug T-Expand /million T cells was selected for future use.

Next, the efficacy of T-Expand for engineering T cells with a CAR was evaluated. Activation with T-Expand prior viral transduction resulted in a T cell transduction efficacy of ∼50% CAR expressing T cells when infected with a multiplicity of infection (MOI) of 3 (MOI-3), which was similar to levels obtained with Dynabeads™ (**Figure 3f**). In contrast, activation with TransAct™ resulted in 20% transduction efficiency (**Figure 3f**) and an overall lower yield of T cells **(Supplementary Figure 14a**) after 10 days of expansion (**Figure 3g**). Phenotypic analysis revealed that T-Expand CAR T cells had a significantly higher percentage of CD27+CD28+ CAR T cells (**Figure 3 h, i, j**) and higher percentage of TEM (**Figure 3k, Supplementary Figure 14b**) with an increased CD8+ CAR T cell population compared to Dynabeads^TM^ and TransAct^TM^ (**Supplementary** Figure 14c, d). For further functional analysis, we compared T-Expand against Dynabeads™ expanded CAR T cells.

To determine the proliferative potential of anti-CD19 CAR T cells after expansion with T-Expand, expanded and cryopreserved CAR T cells were stained with CellTrace Violet and stimulated with irradiated CD19+ Jeko-1 cells to assess their expansion potential upon target recognition (**Figure 4a**). On day 6, flow cytometry analysis showed that ∼85% of T-Expand expanded CD19 CAR T cells had undergone up to four rounds of population doublings, in comparison to only 20% of Dynabeads™-expanded CAR T cells (**Figure 4b**). To determine the cytotoxic potential of CAR T cells expanded with T-Expand, we performed an incucyte based cytotoxic assay with co-cultivation of CAR T cells and Jeko-1 cells at effector:target ratio (E/T) 5:1; 1:1; 1:5; 1:10. **(Figure 4c).** CAR T cells expanded by T-Expand exhibited potent cytotoxicity against CD19+ Jeko-1 cells. Compared to CAR T cells expanded with Dynabeads™, CAR T cells expanded with T-Expand showed significant tumor cell-killing efficacy across all E/T ratios. To further investigate the serial cytotoxicity and phenotypic effect on CAR T cells after serial exposure to target cells, we performed a re-challenge killing assay, by continuedly exposing remaining CAR T cells with CD19 target cells every 2-3 days. Based on the E/T ratio at 5:1, T-Expand-expanded CAR T cells exhibited significantly higher killing capacity even after three rounds of re-challenge (**Figure 4 d, e, Supplementary Fig 15**).

**Figure 4.**
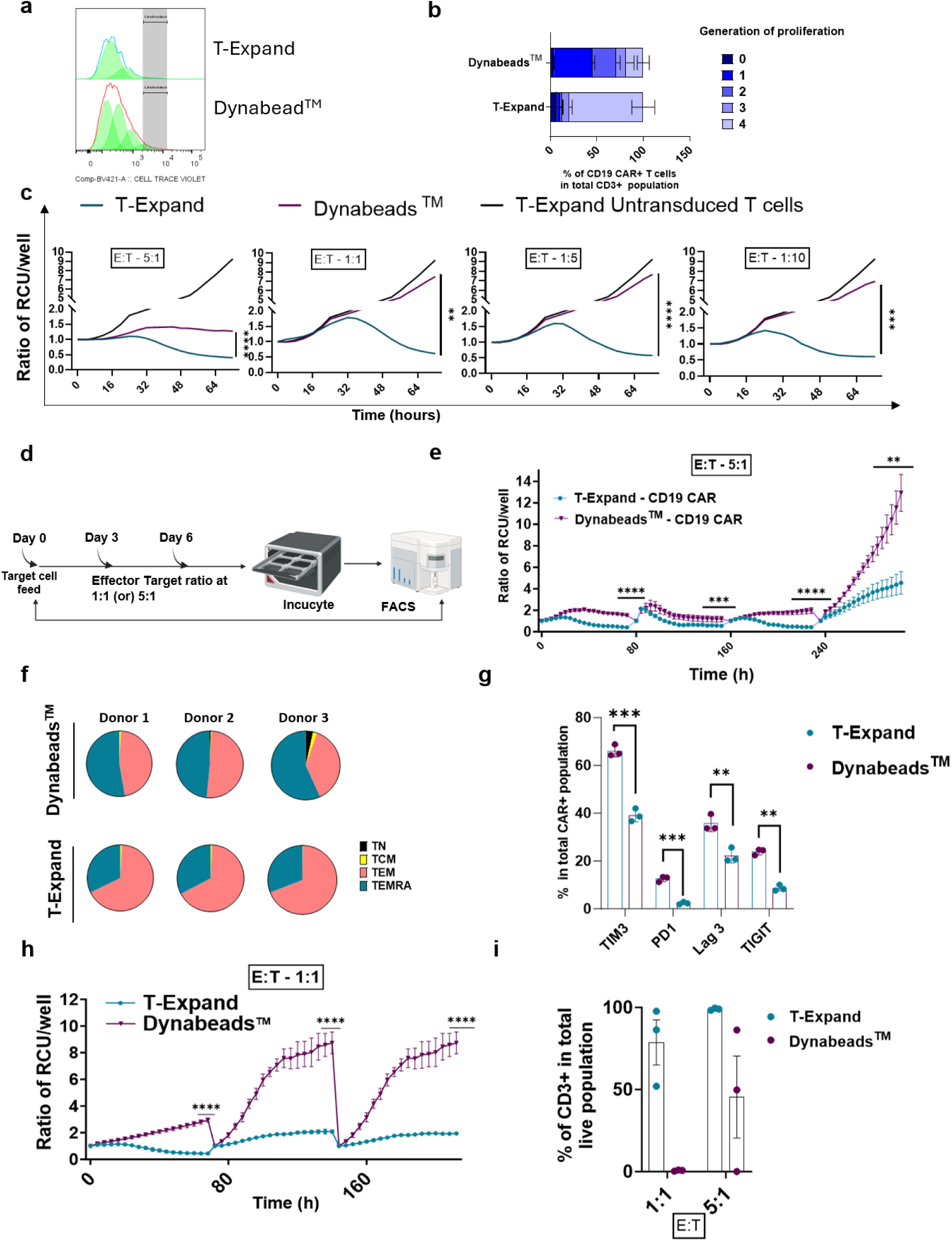
T-Expand enhanced the proliferation and cytotoxic capacity of anti-CD19 CAR T cells. **a,** Representative histograms showing the proliferation peaks on Day 6 of restimulation. Proliferation is determined by flow cytometry analysis (proliferation index in Flowjo by measuring the dilution of cell trace violet. **b,** graph showing the percentage of CAR T cells in different generations of proliferation quantified based on the dilution of cell trace violet and proliferation index analysis. **c,** Cytotoxicity of CAR T cells in co-culture with Jeko-1 cells at an E/T ratio 5:1; 1:1; 1:5; 1:10; tracked over 72 hours in the Incucyte, represented by ratio of relative confluence units (RCU) per well normalized to 0h. **d,** Schematic illustration of the incucyte based cytotoxicity re-challenge assay. **e,** Cytotoxicity of CAR T cells in co-culture with Jeko-1 cells at an E/T ratio of 5:1, tracked over 240 hours in the Incucyte with re-challenge every 48 hours, represented by ratio of relative confluence units (RCU) per well normalized to 0h. **f,** Pie chart showing the percentage of CAR T cells with a CD45RA+CCR7- terminally differentiated phenotype and CD45RA-CCR7- effector memory phenotype. **g,** Graph showing the percentage of CAR T cells expressing PD-1; Tim3; LAG3 and TIGIT surface markers after 3 rounds of rechallenge with the Jeko-1 target cells. **h,** Cytotoxicity of CAR T cells in co-culture with Jeko-1 cells at an E/T ratio of 1:1, tracked over 200 hours in the Incucyte with re-challenge every 48 hours. **i,** Bar graph showing the percentage of CD3+ T cells in the total live cell population at E/T ratio of 1:1 and 5:1 after three rounds of re-challenge. *Data in panels **c**, **e**, **h** represent pooled results from three biological replicates presented as mean ± SD, statistical significance determined by paired t test and **i** represent pooled results from one experiment, presented as mean ± SD, with each dot indicating one PBMC sample. Statistical significance determined via unpaired t-test: *P < 0.05, **P < 0.01, ***P < 0.001, ****P < 0.0001. Data panel in **b** represent results from three replicates presented as mean with ± SD, Statistical significance determined via one-way Anova: *P < 0.05, **P < 0.01, ***P < 0.001, ****P < 0.0001*.

To further characterize CAR T cells with enhanced cytotoxic capacity after repetitive stimulation, remaining cells from the third re-challenge were harvested and analyzed by flow cytometry (**Supplementary Fig 15a**). T cell expanded with Dynabeads™ primarily had a terminally differentiated phenotype whereas CAR T cells expanded with T-Expand were primarily effector memory (**Figure 4 f, Supplementary Fig 15b**). Additionally, the percentage of the CAR T cells expressing exhaustion markers such as PD1, LAG3, TIM3, and TIGIT were all significantly lower in T-Expand compared with Dynabead™ expanded CAR T cells (**Figure 4 g, Supplementary Fig 15c**). Next, we investigated the cytotoxicity of CAR T cells expanded using T-Expand under low E/T ratios. Similarly, CAR T cells expanded using T-Expand demonstrated significantly higher cytotoxicity, even after three rounds of re-challenge (**Figure 4h**). Flow cytometry analysis revealed that under co-culture conditions with E/T ratios of 1:1 and 5:1, the T-Expand group exhibited T cell persistence rates of approximately 70% and 99%, respectively, whereas the Dynabeads™ group showed only 0.2% and 45%. (**Figure 4i, Supplementary Fig 15 d**). This suggests that the superior cytotoxic capacity of T-Expand CAR T cells is likely attributed to a combination of enhanced proliferative capacity, effector memory phenotype and lower exhaustive profile.

### T-Expand expanded CAR T cells show superior *in vivo* tumor killing efficacy

Subsequently, we tested the *in vivo* antitumor efficacy of the CD19 CAR T cells expanded with T-Expand in NXG mice injected with the lymphoma cell line Jeko-1 engineered to express luciferase (luc-mCherry+) for monitoring tumor burden using bioluminescence (BLI). Initially we tested a dose of 1 × 10^6^ CAR T cells expanded with either T-Expand or Dynabeads™ (**Figure 5a**). The tumor growth measured by BLI shows that, the treatment with T-Expand expanded CAR T cells lead to complete tumor elimination by day 7 without signs of tumor relapse at study termination on day 21. However, the Dynabead™ expanded CAR T cell treated group showed insufficient tumor clearance on day 7, further leading to the gradual increase in tumor burden until day 21. (**Figure 5b, c, Supplementary Figure 16**). At study termination, mice were euthanized and spleen and bone marrow were collected for detection of CAR T cells. The results show that the percentage of the CAR T cells detected both in bone marrow and spleen is similar between the T-Expand and the Dynabeads^TM^ group **(Figure 5d)**. Interestingly, mice treated with T-Expand expanded CAR T cells had a significantly higher proportion of CD4+ T cells in the spleen **(Figure 5e)** and higher levels of CD137 (4-1BB) in bone marrow compared to Dynabeads™ expanded CAR T cells **(Figure 5f).** CD137 expression on T cells serves as a marker for antigen specific activation often correlated with enhanced anti-tumor efficacy. CD137 receptor interaction with 4-1BB ligand also plays a significant role in enhancing the survival and proliferation of these T cells(*19*). Next, to further elucidate differences between CAR T cells expanded with T-Expand or Dynabeads™, mice were injected with 1 × 10^6^ luc-mCherry+ Jeko-1 cells and treated on day 7 with 0.25 × 10^6^, 0.5 × 10^6^ or 1 × 10^6^ CAR T cells expanded with either Dynabeads™ or T-Expand (**Figure 5g**). On day 21, blood was collected for further analysis. A higher percentage of circulating T cells in mice receiving CAR T cells expanded with T-Expand (**Supplementary Figure 17a**). The mice treated with T-Expand expanded CAR T cells at lowest doses (0.25 × 10^6^ and 0.5 × 10^6^) showed some tumor control and the high dose (1 × 10^6^) completely eradicated the tumors(**Figure 5h**). In contrary, Dynabeads™ expanded CAR T cells at all doses failed to show any significant tumor control; most mice developed systemic hematological tumors until day 33. (**Figure 5i, Supplementary Figure 17b**).

**Figure 5.**
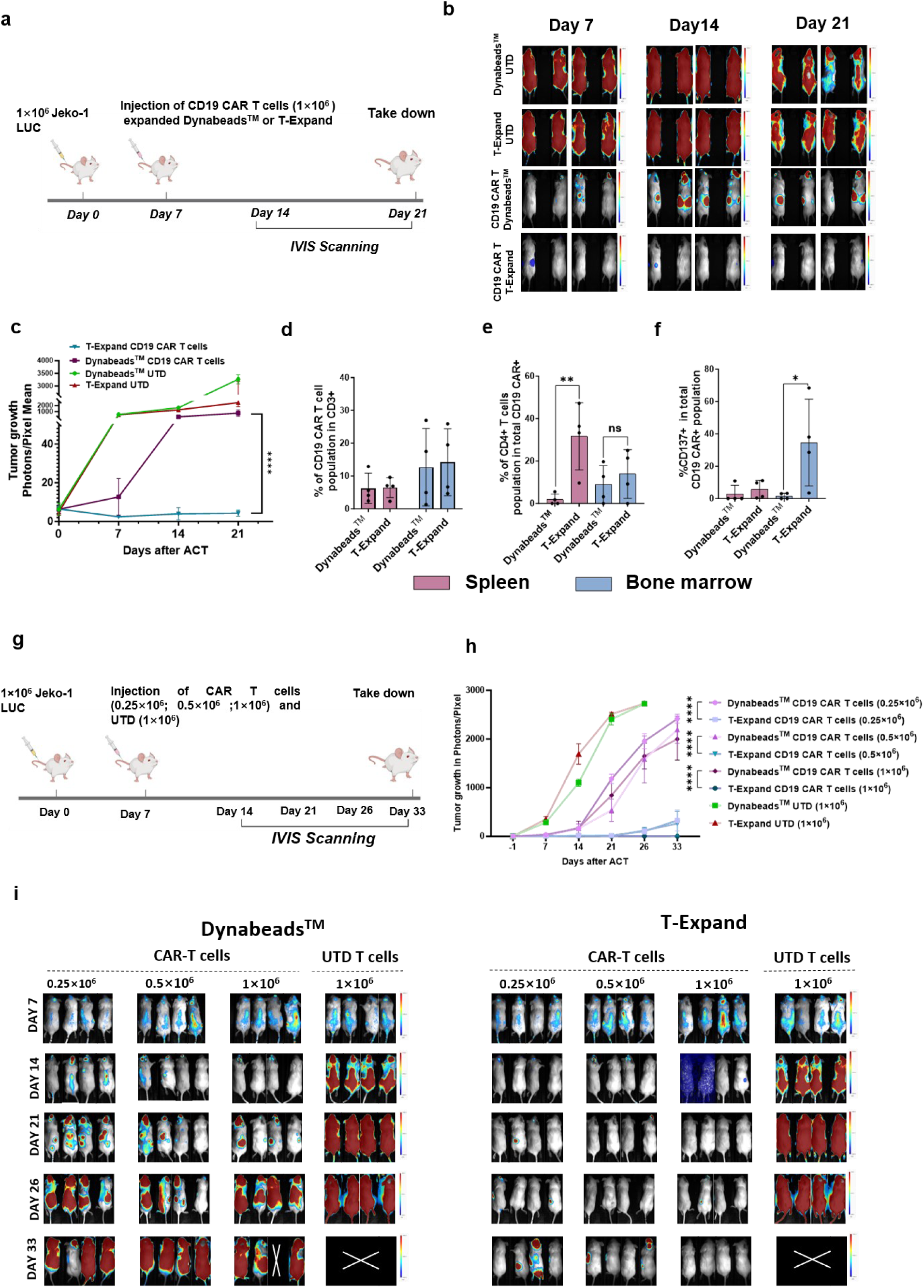
T-expand expanded CAR T display superior antitumor activity in a xenograft model of B-cell lymphoma. **a,** Schematic of the *in vivo* study setup. 1×10^6^ Jeko-1 lymphoma cells expressing mCherry and luciferase were injected into NXG mice. Tumor volumes were measured on day 7, and mice were randomized by tumor burden, receiving either T-Expand or Dynabeads™ expanded CAR T cells (mentioned dosage) or UTD T cells at the highest CAR T cell dose equivalent. Bioluminescence from luciferase-expressing tumor cells was tracked using IVIS imaging every 7 days. **b,** Representative IVIS overlay images showing bioluminescent signals for each treatment group. **c,** Bioluminescence intensity, reported as photons/pixel, tracked over time for each treatment group. The mice were taken down on day 21 after ACT and the spleen and bone marrow were analyzed in flow cytometry. **d,** Graph showing % of CD19 CAR T cells in the CD3+ T cell population in spleen and bone marrow. **e,** Graph showing % of CD4+ T cells in the CAR+ T cell population in spleen and bone marrow. **f,** Graph showing % of CD137+ T cells in the CAR+ T cell population in spleen and bone marrow. **g,** 1×10^6^ Jeko-1 lymphoma cells expressing mCherry and luciferase were injected into 32 NXG mice. Tumor volumes were measured on day 7, and mice were randomized by tumor burden, receiving either CAR T cells at various doses or UTD-T cells at the highest CAR T cell dose equivalent. Bioluminescence from luciferase-expressing tumor cells was tracked using IVIS imaging every 7–10 days. **h,** Bioluminescence intensity, reported as photons/pixel, tracked over time for each treatment group with each line representing a mouse. **i,** Representative IVIS overlay images showing bioluminescent signals for each treatment group; *Data in panels **e**, **f** represent pooled results from 1 experiment, presented as mean ± SD with every dot (or) line representing a mouse. Statistical significance determined via unpaired t-test: *P < 0.05, **P < 0.01, ***P < 0.001, ****P < 0.0001* *Data in panels **c** and **h** represent pooled results as mean ± SD with 4 mice in each group. Statistical significance determined via one-way Anova with Šídák’s multiple comparisons test on day 33 of CAR T cell injection of each group: *P < 0.05, **P < 0.01, ***P < 0.001, ****P < 0.0001*

### Deep RNA sequencing analysis of expanded CAR T cells

To investigate the transcriptomics of expanded CAR-T cells, we collected CAR T cells on Day 10 of expansion using either T-Expand or Dynabeads^TM^ for bulk RNA sequencing from four healthy donors. The principal component analysis (PCA) revealed that the overall gene profiles of T-Expand expanded CAR T cells markedly differed from those of Dynabeads^TM^ expanded CAR T cells (**Supplementary Figure 18a, b**). T-Expand expanded CAR T cells exhibited high expression of genes associated with a putative transitional state between cytotoxic effector and differentiation phenotypes (**Figure 6a**). These transitional phenotypes were identified by the co-expression of multiple cytotoxic genes, including LTB and GZMA(*20*).

**Figure 6.**
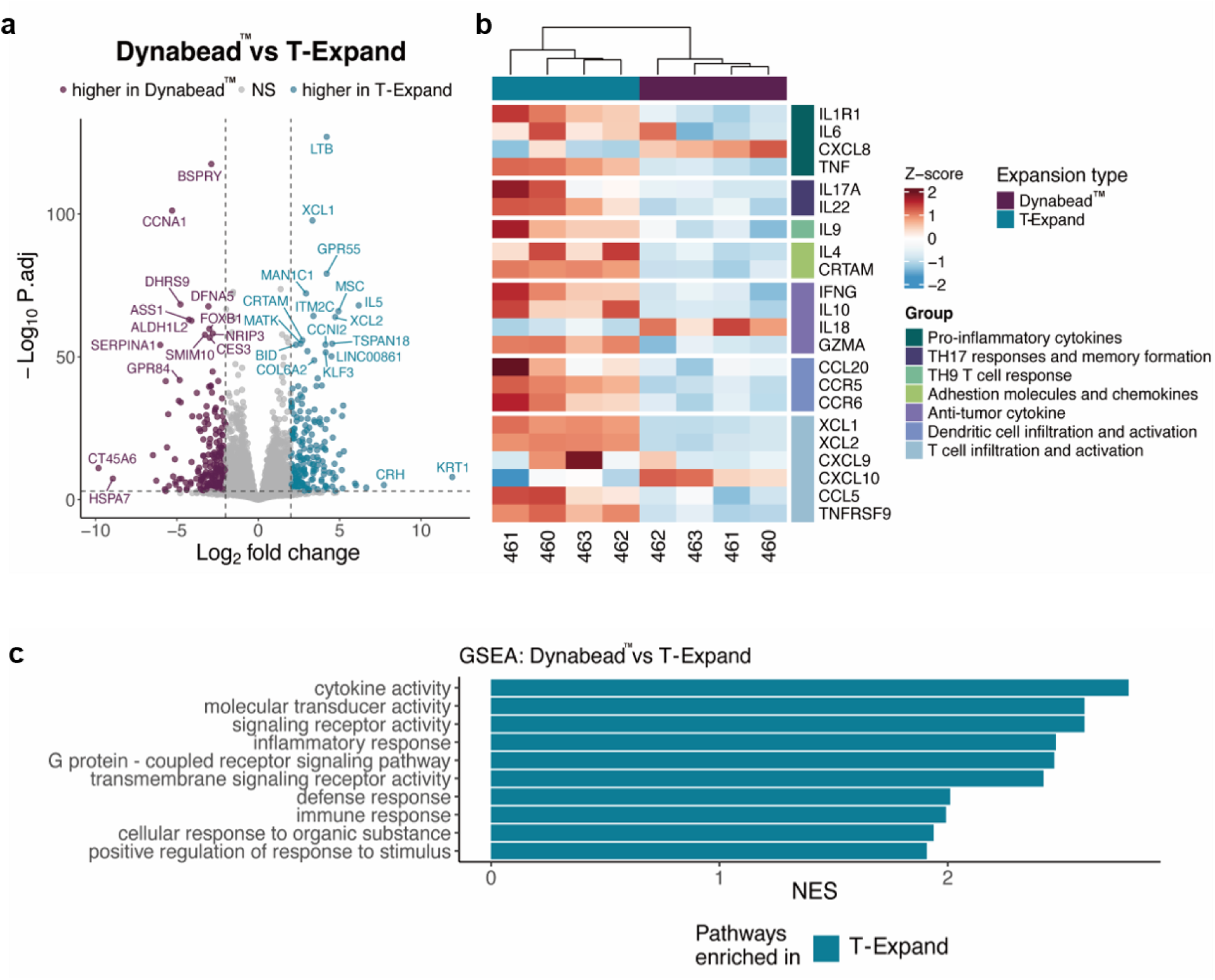
Differential gene expression in CAR T cells expanded with either T-Expand or Dynabeads^TM^, highlighting key changes between the two expansion methods. **a,** Volcano plot of differentially expressed genes between T-Expand and Dynabeads^TM^. **b,** Heatmap showing selected genes grouped based on known functions. **c,** GSEA was performed on bulk RNA-seq data. Normalized Enrichment Score (NES) plot of the top 10 gene sets found in T-Expand. To keep only highly significant gene sets, we used only gene sets with >30 genes and with p-values < 0.001. Using these strict thresholds, all gene sets enriched in Dynabeads^TM^ disappeared.

The analysis showed that the T-Expand expanded CAR T cells also showed upregulation of Xcl1 & Xcl2 genes that are involved in dendritic cell recruitment and activation. The stem-like CTLs express high levels of Xcl1 that promotes the infiltration of migratory cDC1s that are associated with improving CD8 T cell anti-tumor activity through cross-primin(*21, 22*). Among the selected gene set (**Figure 6b**), CAR T cells expanded with T-Expand showed increased expression of genes associated with anti-tumor cytokines and effector molecules (IFNg, GZMA), as well as T cell infiltration and activation (CXCL9, CXCL10, CCL5), compared with Dynabeads^TM^ expanded CAR T cells(*23*),(*24*),(*25*). (**Supplementary Fig 18 b**)

To further elucidate the global transcriptomic landscape related to T-cell functionality in T-Expand expanded CD19 CAR T cells, only significantly expressed genes (p.adj < 0.001) were selected, and the top 10 gene sets were characterized via gene set enrichment analysis (GSEA) (**Figure 6c**). The results revealed a significant enrichment of gene sets related to immune responses, inflammatory regulation, and signal transduction in T-Expand expanded CAR T cells. In contrast, Dynabeads^TM^ expanded CAR T cells did not show notable enrichment of these gene sets, indicating substantial differences in gene expression regulation between these two expansion strategies.

## DISCUSSION

In this study, we successfully developed T-Expand, a novel nanoplatform designed for the efficient expansion of T cells. Compared with existing T cell expansion platforms (Supplementary Table 1), T-Expand excelled in promoting T cell activation and proliferation, outperforming previously reported nanoplatforms and even surpassing certain micron-scale aAPC platforms. By functionalizing native dextran, we introduced high-density azido groups through click chemistry on its abundant modifiable side chains, thereby enabling the conjugation of anti-CD3 and anti-CD28 antibodies and effectively mitigating the low expansion efficiency caused by insufficient antibody density(*26*). We optimized the stoichiometric ratio of anti-CD3 and anti-CD28 antibodies, achieving both robust T cell yields and favorable phenotypic characteristics of the expanded T cell products. Furthermore, the T-Expand’s excellent biocompatibility and nanoscale advantages eliminate the need for additional removal steps following T-cell expansion.

we employed genetically engineered Jurkat cells expressing mCherry-Cathepsin L to visualize how T-Expand induces Jurkat cell activation. Leveraging the multifunctionality of T-Expand, we loaded BV421-streptavidin into T-Expand to track its interactions with T cells. Once naïve T lymphocytes are activated, CD4(+) T cells—similar to CD8(+) T cells—polarize labeled lysosomal granules toward their target cells(*27*). Based on this observation, when T-Expand interacts with Jurkat cells, the uniformly distributed mCherry-labeled Cathepsin L granules co-localize and form prominent clusters around T-Expand. Comparative studies further showed that, relative to Dynabeads™-activated cells, Jurkat cells activated by T-Expand exhibit significantly enhanced multicentric granule trafficking and aggregation. These findings suggest that T-Expand is capable of activating Jurkat cells at multiple sites and effectively inducing granule polarization toward the T-Expand contact regions.

Moreover, activation with T-Expand successfully lead to virally transduced CAR expression in approximately 50% of T cells, achieving similar results as Dynabeads™ and outperformed TransAct™. Our results demonstrate that T-Expand-expanded CAR T cells exhibit a less differentiated phenotype, featuring a high proportion of CD27+CD28+ effector memory T cells, compared to those generated using conventional platforms such as Dynabeads™ and TransAct^TM^, which predominantly produced TEMRA cells. This indicates that T-Expand facilitates the generation of CAR T cells with a potentially more favorable phenotype for therapeutic efficacy.

In cytotoxicity assays, CAR T cells expanded by T-Expand demonstrated outstanding anti-tumor activity. The CD19 CAR T cells maintained high viability and consistent killing activity after three rounds of re-challenge with CD19+ Jeko-1 target cells. Phenotypic analysis post-rechallenge of the CAR T cells revealed a significantly increased proportion of effector memory T cells and reduced expression of exhaustion markers, indicating that CAR T cells possess enhanced durability and survival potential in anti-tumor responses. This optimal cellular profile is likely key to the superior functionality and persistence of T-Expand expanded CAR T cells as observed in the *in vivo* studies. Based on the lymphoma model, a dose of 0.25 × 10^6^ CAR T cells expanded with T-Expand achieved robust anti-tumor efficacy, even outperforming the high-dose treatment group of 1 × 10^6^ CAR T cells produced with Dynabeads™. This remarkable tumor-killing efficacy in vivo aligns with our in vitro cytotoxicity and repetitive restimulation assays. CAR T cell products with a younger effector memory phenotype (CD27⁺ and CD28⁺) and heightened proliferation and cytotoxic capacity have demonstrated improved prognoses in clinical applications(*28–32*).

Bulk RNA sequencing of CAR T cells expanded using the T-Expand platform revealed significant upregulation of key effector molecules (GZMA, EOMES, IFN-γ, TNFRSF9/CD137), aligning with their sustained cytotoxicity and robust rechallenge capacity in vitro and in vivo. These cells also showed elevated expression of Th2 (IL4, IL5, IL10, IL13), Th9 (IL9), and Th17 (IL17A, IL17F, IL22) cytokine genes. Recent studies suggest that Th2 CAR T cell products provide superior expansion and antitumor efficacy—particularly after leukemia relapse— and that enhancing Th2 functionality can partially restore antitumor effects.(*33–35*). T-Expand expanded CAR T cell treated mice showed significant increase in the proportion of CD4⁺ T cells in the spleens. Additionally, reports have highlighted that Th17 CAR T cells are resistant to cellular senescence^34^. Notably, T-Expand–expanded CAR T cells also showed high expression of multiple chemokine receptor genes (CCR1, CCR2, CCR3, CCR5, CCR6, CCR10) with the high expression of chemokines (CCL1, CCL3, CCL4, CCL20), which may facilitate the recruitment of other immune cells, further amplifying and modulating local immune responses as well as enhancing CAR-T cell infiltration into tumors microenviroment(*36*). However, we couldn’t confirm these compartments of the anti-tumor activity due to the immunodeficient nature of our NxG mouse model. Overall, bulk RNA sequencing demonstrates significant gene expression differences between CAR T cells expanded using the T-Expand and those using Dynabeads^TM^. These molecular features provide potential mechanistic support for the superior cytotoxicity and sustained anti-tumor effects observed with T-Expand expanded CAR T cells.

In summary, T-Expand has some properties that are beneficial to T cells in the context of adoptive cell therapy. T-Expand not only efficiently produces a clinically relevant proportion of CAR T cells, but also creates a more beneficial T cell subset that is associated with enhanced proliferation, persistence, and anti-tumor efficacy by modulating cell phenotype and reducing exhaustion. Thus, T-Expand offers a novel approach to ensuring the production of high-quality CAR T cells and underscores the potential of T-Expand as a promising alternative to traditional activation strategies.

## Methods

### Preparation of dextran NPs (naked NPs)

In general, acetalated azido modified Dextran (20 mg) was dissolved in cold dichloromethane (DCM, 500 μL). The PBS (100 μL) was added to undergo a sonication for 20 s every 10 seconds on ice using a probe sonicator 9 (Qsonica L.L.C, USA) flat tip, an output setting power of 5. Then the poly(vinyl alcohol) (PVA, Mw 9000-10000 g/mol, 80% hydrolyzed) (1 mL, 3% w/w in MQ water) was added into the primary W/O solution and further sonicated for an additional 20 s on ice using the same settings. The resulting W/O/W emulsion was immediately transferred to the a PVA solution (5 mL, 0.3% w/w in MQ water) and stirred for 3 h at room temperature for evaporating the DCM solvent. The NPs were obtained by centrifugation (10.6 g, 10 min) and washed with PBS 3 times.

### Formation of DBCO functionalized antibody

First, the anti-CD3 and anti-CD28 antibodies were dissolved in PBS buffer (0.5 mg/ml), and the DBCO-Sulfo-NHS was dissolved in DMSO to final concentration at 10 mM. Next, 3.33 μL of the above DBCO-Sulfo-NHS Ester was added to each antibody solution and incubated at room temperature for 30 minutes. Then, the reaction was quenched with 18 ul of Tris-buffer (1 M, pH 8) for 15 minutes. Finally, any unbound DBCO-Sulfo-NHS Ester was removed using Zeba spin desalting columns (7 kDa or 40 kDa MWCO, Thermo Fisher Scientific).

### Formation of T-Expand for T cell activation and expansion

Generally, DBCO-functionalized and labeled antibodies (anti-CD3-DBCO and anti-CD28-DBCO) were conjugated onto the surface of azido modified dextran NPs via click chemistry. The initial concentration of reacting antibodies was varied with various antibody ratio prepared. After the incubation in room temperature for 2 hours, the T-Expand was obtained by centrifugation (10.6g, 10 mins) and stored in 4-degree fridge for further using.

### Nanoparticle characterization

The number and size of the NPs was measured by Nanoparticle Tracking Analysis (NTA, NANOSIGHT, Malvern Instruments). The aliquot of NPs solution was diluted in MiliQ water pre-filtered by 20 nm filter at the same numbers used for expansion. The setup for NTA test was shown as follows (Camera Type: sCMOS Laser Type: Green; Camera Level: 10; Slider Shutter: 696; Slider Gain: 55; FPS 25.0; Number of Frames: 1498; Temperature: 23.9 °C; Viscosity: (Water) 0.910 - 0.911 cP; Syringe Pump Speed: 100).

### Cryo-Transmission electron microscopy

Three µL of the sample was applied on a hydrophilized lacey carbon 300 mesh copper grid (Ted Pella Inc., California, USA). The excess sample on the grid was blotted with filter paper at blotting time of 3 s, blotting force 0, temperature 4°C, and 100% humidity (FEI Vitrobot IV, Eindhoven, The Netherlands), and was rapidly plunged into liquid-nitrogen cooled ethane (- 180⁰ C). Sample observations were performed using a Tecnai G2 20 transmission electron microscope (FEI, Eindhoven, The Netherlands) at a voltage of 200 kV under a low-dose rate. Images were recorded with an FEI Eagle camera 4kx4k at a nominal magnification of 29 kX.

### Quantifying the Azido molar on the surface of Dextran nanoparticles

The sulfo-Cyanine5 DBCO (1mg) was dissolved in 1 mL of DMSO. The concentration calibration curve was obtained by detecting the servals concentration (0.25 ug/ml, 0.5 ug/ml, 0.75 ug/ml, 1 ug/ml, 2.5 ug/ml) of sulfo-Cyanine5 DBCO. Next, 10 ul of dextran NPs were incubated with 1 ul of sulfo-Cyanine5 DBCO (1 mg/ml) overnight. The NPs were pelleted at the bottom of the centrifuge tube by centrifugation at 10600 g for 10 minutes. The supernatant of un-clicked sulfo-Cyanine5 DBCO was measured by plate reader. The amount of clicked sulfo-Cyanine5 DBCO was determined by calculating the concentration difference before and after coupling.

### Nanodrop based quantification of antibody conjugation

The DBCO-modified antibody was dissolved in 600 µL of PBS solution, and its concentration was measured using a NanoDrop spectrophotometer. Next, dextran nanoparticles or T expand particles, intended for T cell expansion and CD19 CAR T cell manufacturing, were added to the antibody solution. After incubating the mixture overnight, the nanoparticles were pelleted at the bottom of the centrifuge tube by centrifugation at 10600 g for 6 minutes. Subsequently, the supernatant was then collected, and the antibody concentration was measured again with the NanoDrop. The amount of conjugated antibody was determined by calculating the concentration difference before and after coupling.

### Primary T-cell isolation and activation

Peripheral blood mononuclear cells (PBMCs) were sourced from healthy donors from Rigshospitalet, Copenhagen as approved by the Danish ethics committee. PBMCs were isolated from whole blood by density centrifugation using SepMate tubes (StemCell) and frozen at -80°C in FBS and 10 % DMSO.

Primary T cells from healthy donor PBMCs were isolated using the EasySep™ Human T cell enrichment kit supplied by STEMCELL technologies (as per supplier instructions). The purified T cells were resuspended in 0.5 × 10⁶ cells/ml in RPMI+10%FBS+1% Penstrep (supplied by Thermo Fischer Scientific) and IL2 (20IU/µl) (supplied by PeproTech®). For T cells activation, 100,000 purified T cells were plated in 100 μL culture medium in a 96-well cell culture plate, then specific concentration (0.75 mg/ml: 1ul, 2ul, 5ul, 10 ul) of T-Expand (or) Dynabeads^TM^ at a bead-to-cell ratio of 3:1 (or) TransAct^TM^ were added to each well and incubated in a humidified CO_2_ incubator at 37°C for 20 hours. Next, the activated T cells were harvested and washed with PBS buffer for further flow cytometry analysis.The T cell activation is measured by staining the T cell activation markers CD69 and CD25. The expression levels of these CD69 and CD25 markers can be measured by detecting the number of fluorochrome-labelled cells using LSRFortessa™ flow cytometer (BD Biosciences, USA) as a percentage of the

For T cells expansion, 1,000,000 purified T cells/mL and IL2 (20 IU/µL) in a 48 well cell culture plate or suitable tissue culture plate or tissue culture flask. Then added 20 μL or 40 ul of T-Expand (0.75 mg/ml) (or) Dynabeads^TM^ at a bead-to-cell ratio of 3:1 (or) TransAct^TM^ and incubated in a humidified CO_2_ incubator at 37°C.

The T cells were counted on day 3, 5, and 7, and their viability was reassessed. The cells are resuspended in fresh media to maintain a concentration of 1 × 10^6^ cells/mL, and rhIL-2 is replenished to a final concentration of 20 IU/mL to support continued growth and expansion. On Day 10, the culture contained Dynabeads^TM^ was thoroughly resuspended to ensure uniform mixing of cells and beads which was then removed using a magnetic separator, while the T-Expand expanded CAR T cells at MOI of 3 were washed with fresh cell medium and further transferred to a clean container. Finally, the cells are cryopreserved for downstream application.

### Cell lines and culture

The human cell lines Jeko-1(B cell lymphoma) obtained from American Type Culture Collection (ATCC) and cultured in RPMI supplemented with 10% fetal bovine serum and 1% penniciln streptomycin. For cytotoxicity assay and in-vivo tumor models, these cell lines were transduced with mCherry fluorescent protein and luciferase genes using lentiviral vectors and were sorted on a FACS melody (BD Biosciences, USA) to establish new cultures.

### Flow cytometry

Cells were stained with antibodies against phenotypic, intracellular markers, run on a LSRFortessa™ flow cytometer and the data analyzed using the FlowJo software. Cell lines were sorted on a FACSAria™ flow cytometer (BD Biosciences, USA) using a 100 µm nozzle. Amplitude was adjusted to optimize droplet break-off, and droplet calibration was performed before each sort using AccuDrop™ beads (BD, USA).

### Airyscan microscope imaging of T-Expand interacting with Jurkat cells

The engineered Jurkat cells were cultured with T-Expand or Dynabeads^TM^ for 24 hours. Following incubation, centrifugation was conducted to remove the culture medium and resuspend the cells in FACS buffer (PBS + 1% BSA). Cell membrane was stained with anti-human CD45 and incubated on ice for 20 min. The cells were subsequently collected by centrifugation and 5 µl of the cell suspension was transferred to a glass slide (Thermo scientific). The cell droplet was mounted with a coverslip and the edges sealed with nail polish. Confocal images were obtained by using Airyscan confocal microscope (LSM 900 Airyscan 2, Carl Zeiss Microscopy GmbH). For real-time imaging, engineered Jurkat cells were cultured in an 8-well Glass µ-slide chamber (ibidi) at 37°C and stimulated with T-Expand or Dynabeads^TM^ for 24 h. 3D Z-stack imaging was conducted using an Airyscan laser confocal microscope, with subsequent analysis performed using Imaris software (v.10.1.0).

### CAR construct and Lentiviral production

A second-generation CAR were designed with an anti-CD19 (FMC63) CD8TM/hinge domain, 4-1bb, and CD3ζ. Expression of the CAR was linked with expression of a green fluorescent protein (GFP) 27 reporter gene. The CAR was inserted into a third-generation lentiviral vector under the control of a human EF1α promoter. All plasmids were synthesised by GenScript, USA. Lentiviral particles were generated in HEK-293T cells by using LipofectamineTM for transfecting them with three separate packaging plasmids: pMDLg/pRRE was a gift from Didier Trono, pRSV.REV (Addgene plasmid #12253 https://www.addgene.org/12253/)36, pMDLg/p.RRE (Addgene plasmid #12251 https://www.addgene.org/12251/), and pMD2.G (Addgene plasmid #12259 https://www.addgene.org/12259/), along with a transfer plasmid containing the DNA construct of interest(*37*). The transfected HEK-293T cells are incubated at 37°C and 5% CO2 and the cell culture supernatant were collected at 24 and 48hrs time point. These supernatants were up concentrated for lentiviral particles using Lenti-X™ Concentrator according to the maufacturer’s recommendations (Takara bio).

The titter of the produced lentivirus particles was assessed through a titration experiment by infecting SUP-T1 cells and subsequently analysing GFP or surface expression of the receptor via tetramer staining via flow cytometry.

### Engineering of Jurkat with mCherry-Cathpesin L

A cathepsin L-mCherry fragment was synthesized and cloned into the same backbone as the CD19 CAR by GenScript, USA. Lentivirus was created following the same protocol as the previous setup, and Jurkats were transduced using an MOI of 5, followed by sorting of the cells based on mCherry expression.

### Production of CAR T cells *in vitro*

On Day 0, the purified human T cells from healthy donors were diluted to 1 × 10⁶ cells/mL in RPMI media containing recombinant human IL-2 (rhIL-2) at a final concentration of 20 IU/mL, which was then cultured with T-Expand (0.75 mg/ml, 20 ul), or TransAct^TM^ (10 ul) or Dynabeads^TM^ (Gibco™ Anti-CD3/CD28 Dynabeads™) based on a bead-to-cell ratio, typically 3:1 or 1:1 for 24 hours. Then the CD19 CAR lentiviral particles were added to the culture at a multiplicity of infection (MOI) of 3 to ensure efficient transduction. Further the CAR T cells are expanded using the T cell expansion protocol.

### In vitro T-cell phenotype analysis

According to the experimental design, the phenotype of T cells and CAR T cells was evaluated using flow cytometry. The transduction efficiency of CAR was determined by the co-expression of two markers: the intracellular fluorescent protein GFP encoded by the CAR construct and CD19 tetramer. Flow cytometry staining was performed on ice for 20 minutes in FACS buffer. The concentration of antibodies used for staining was determined according to the manufacturer’s instructions. Detailed information on the antibodies is provided in the **Supplementary Table 3**. Stained cells were analyzed using an LSRFortessa flow cytometer, and data analysis was conducted using FlowJo v10 software. Gates were set for each time point and sample independently based on fluorescence minus one (FMO) control.

Cells were centrifuged at 500 x g for 5 minutes at 4°C, and the supernatant was discarded. The pelleted cells were resuspended in 5 µL of 1 µM Dasatinib (LC Laboratories, D3307), antigen tetramers were added, and the cells were incubated for 15 min at 37°C in the dark. The cells were washed once with FACS buffer, followed by staining with Near-IR 28 (NiR) viability dye (Invitrogen™, L34976) and additional antibodies for surface staining for surface markers 30 min at 4°C in the dark. Finally, the cells were washed twice with FACS buffer in 200-300 µL of FACS buffer, and immediately analysed on a LSRFortessa™ flow cytometer. Alternatively, the cells were fixed with 50 µL of 1% paraformaldehyde for 1-2 h (if required), washed twice with FACS buffer, and analyzed 2-24 hours late

### Proliferation assay

The T cells were resuspended 1 × 10⁶ cells/mL in PBS and 0,75ul/ml of CellTrace violet is added to the T cells and incubated at 37°C for 15mins. After incubation and co-cultured with irradiated Jeko-1 target cells at a 1:5 effector-to-target ratio for 6 days. Flow cytometry was used to measure CellTrace Violet dilution, and the proliferation index was calculated using FlowJo software.

### In vitro CAR T cell killing assays and rechallenge assay

The CD19 CAR T cells and the CD19 target cells, Jeko 1 lymphoma cell line modified to express mCherry, resuspended in RPMI supplemented with 10%FBS and 1% Pen/Strep are seeded at the mentioned effector:target ratio, for eg., 1:1. On Day 0, 1 × 10⁴ target cells and 1 × 10⁴ CAR T cells were co-cultured in a 96-well plate with 200 μL of T-cell culture medium. After a 48-hour incubation, the supernatant was collected for ELISA test, followed by the addition of another 1 × 10⁴ target cells to each well. This process was repeated every 48 hours to simulate the repeated in vivo challenge faced by CAR T cells.

### Efficacy studies in vivo

For *in vivo* studies, 6-week-old female NXG Immunodeficient mice (NOD-Prkdcscid-IL2rgTm1/Rj) were acquired from Janvier Labs and housed at the Bio Facility, Department of Health Technology, Technical University of Denmark. All procedures were approved by the Danish National Animal Experiment Inspectorate and the institutional ethical board.

For efficacy studies, 1 × 10⁶ Jeko-1 lymphoma cells expressing luciferase were injected intravenously into 7–10-week-old NXG mice. Tumors were allowed to engraft for 7 days before randomization and treatment start. On treatment day, indicated amount of untransduced, Dynabeads^TM^-expanded T cells or T-Expand expanded CD19 CAR T cells in 200 µl PBS were administered i.v. by tail vein injection. Tumor growth was monitored weekly with bioluminescence imaging (Optical Imaging unit, MILabs). Mice received 30 mg/mL of D-Luciferin intraperitoneally and 20 min later bioluminescence was measured. Measurements were analyzed using fixed-size regions of interest using MILabs optical imaging software. Mouse body weight and appearance were monitored throughout the experiment.

At study termination, spleen and bone marrow were harvested for further analysis by flow cytometry. Spleens and bone marrow were passed through 70 µm filters and washed in PBS to create a single-cell suspension. Splenocytes subsequently went through red blood cell lysis before being passed through a 70 µm filter again. Blood was harvested on day 21 from the saphenous vein. Red blood cells were lysed before washing with PBS. Blood, splenocytes and bone marrow cells were surfaced stained for Near-IR 28 (NiR) viability dye and antibodies against surface markers, CD3, CD4, CD8, CD19, CD137, CD69 and CD19 CAR tetramer and run on LSRFortessa™ for flow cytometry analysis.

### Differential gene expression analysis and GSEA

Differential gene expression analysis was used to identify differentially expressed genes in PBMCs of four donors expanded with either T-Expand or Dynabeads^TM^. Raw RNA-Seq reads were trimmed using Trim Galore and transcript abundances were quantified using Kallisto Quant(*38*). The transcript abundances from Kallisto were imported into DESeq2(*39*) for differential expression analysis. Log2 Fold Change > 2 and < −2 with an adjusted p-value < 0.05 was used as threshold for over- and under-expressed genes for the analysis. The volcano plot was generated with ggplot2 and the heatmap was generated with Complex Heatmap(*40*).

Gene set enrichment analysis (GSEA) was performed using clusterProfiler(*41*) to identify significantly enriched Gene Ontology (GO) terms. As an input for the GSEA we used a ranked gene list of all significant genes (p.adj < 0.001) from the DESeq2 analysis, ordered by Log2 Fold Change. This ranked gene list was then used as input for the gseGO function from the clusterProfiler package to identify significantly enriched Gene Ontology (GO) terms. To retain only highly significant GO terms, this analysis was performed using minGSSize > 30 (selecting only gene sets with more than 30 genes) and p value Cut off < 0.001.

### Statistical analysis

Data are represented as the mean ± standard deviation. When necessary, a two-tailed Student’s *t*-test was used to identify statistically significant differences between the two groups, using GraphPad Prism 8 for computations. In instances of comparison among more than two groups, a one-way analysis of variance (ANOVA) was conducted, followed by Tukey’ s multiple comparison test. Levels of statistical significance were denoted as follows: not significant, *P* > 0.05; \**P* < 0.05; \*\**P* < 0.01; \*\*\**P* < 0.001.

## Author contributions

T. Z., K.R., M.O., Y. S., and S. R. H. designed the experiments. T.Z. contributed to T-Expand design. The *in vitro* and *in vivo* T cell assays were designed and carried out by K.R, T.Z., H.H., M.H., M.O. and C.R.P. The proteomics data was analyzed by H.L. The bulk RNA sequencing data was analyzed by K.K.M. T. Z., K.R. and M.O. contributed to the organization and analysis of the data. T.Z, K.R. and M.O. co-wrote the paper. Y. S. and S. R. H support and supervise this project. All authors discussed the results and help for the revision of the manuscript.

## Acknowledgements

This research was funded in part by: The Novo Nordisk Foundation (Challenge program) (to S. R. H., Y. S. and H. A.). A grant from the European Union’s Horizon 2020 research and innovation program Marie Curie grant agreement No. 955575 (to S. R. H.)

