## supplemental figure 1-18 for "Dextran-based T-cell expansion nanoparticles for manufacturing CAR T cells with augmented efficacy"

Supplementary Fig. 17 | Dose de-escalation experiment to further elucidate differences between CAR T cells expanded with T-Expand or Dynabeads™. 24

Supplementary Fig. 18 | Deep RNA sequencing analysis of expanded CAR T cells expanded with T-Expand or Dynabeads™. 25

**Supplementary tables** 26

Supplementary Table 1. Microscale- and Nanoscale-Based aAPCs for Polyclonal expansion of primary human or mouse T cells 26

Supplementary Table 2. The twenty most abundant proteins in A, B and C samples 28

Supplementary Table 3: antibody and cytokine information 31

**Supplementary references** 32

**Captions for the supplementary videos** 34

**Supplementary methods**

**Chemicals**

Sodium azide (for synthesis), 3-Bromo-1-propanol (97%), dimethylformamide (ACS reagent, ⩾99.8%), ethyl acetate (ACS reagent, ⩾99.5%), magnesium sulfate (anhydrous, ReagentPlus®, ⩾99.5%), Carbonyldiimidazole (CDI, ⩾97.0% (T), for peptide synthesis ), dextran (from Leuconostoc mesenteroides, average mol wt 9,000-11,000), anhydrous dimethyl sulfoxide (DMSO), pyridinium p-toluenesulfonate (98%), 2-methoxypropene (97%), spermine (⩾99.0% (GC)), Dimethyl sulfoxide-d6, poly(vinyl alcohol) (Mw 9,000-10,000, 80% hydrolyzed) were purchased from Sigma-Aldrich.

**Synthesis of azido modified dextran**

***Synthesis of 1-Azido-3-propanol.***

The mixture of sodium azide (0.94 g, 14.4 mmol) and 3-bromo-1-propanol (1 g, 7.2 mmol) was dissolved in 10 mL of dimethylformamide (DMF) and stirred for 24 h at 50 °C. Next, the reaction mixture was filtered out. The DMF was distilled out under reduced pressure. The remaining liquid was extracted by ethyl acetate. The ethyl acetate solution containing product was dried using magnesium sulfate overnight as followed with rotary evaporator to get liquid product.

(Supplementary Fig. 1) $^{1}H$ NMR (400 MHz, DMSO) δ 4.57 (td, J = 5.1, 1.3 Hz, 1H), 3.47 (q, J = 6.2 Hz, 2H), 3.39 (t, J = 6.8 Hz, 2H), 1.68 (p, J = 6.5 Hz, 2H). $^{13}C$ NMR (101 MHz, DMSO) δ 58.09, 48.23, 31.95.

LC-MS (m/z): Calcd for $[M+Na]^{+}$ : 124.1, Found: 124.114

***Synthesis of 8-Azido-3,6-dioxaoctanol.***

The mixture of sodium azide (0.94 g, 14.4 mmol) and 2-[2-(2-Bromoethoxy)ethoxy]ethanol (1.53g, 7.2 mmol) was dissolved in 10 mL of dimethylformamide (DMF) and stirred for 24 h at 50 °C. Next, the reaction mixture was filtered out. The DMF was distilled out under reduced pressure. The rest of the liquid was extracted by ethyl acetate. The ethyl acetate solution containing product was dried using magnesium sulfate overnight as followed with rotary evaporator to get brown liquid product.

(Supplementary Fig. 2) $^{1}H$ NMR (400 MHz, DMSO) δ 4.56 (t, J = 5.4 Hz, 1H), 3.61 (dd, J = 5.7, 4.3 Hz, 2H), 3.56 (ttd, J = 5.7, 3.1, 0.9 Hz, 4H), 3.52 – 3.47 (m, 2H), 3.46 – 3.42 (m, 2H), 3.41 – 3.37 (m, 2H). $^{13}C$ NMR (101 MHz, DMSO) δ 72.84, 70.25, 70.20, 69.73, 60.68, 50.47.

LC-MS (m/z): Calcd for $[M+H]^+$: 176.1; $[M+Na]^+$: 198.1. Found: 176.059, 198.2

***Synthesis of 3-azidopropyl 1H-imidazole-1-carboxylate (Azido-CDI).***

Carbonyldiimidazole (CDI) (1.76 g, 10.85 mmol) was dissolved in 20 mL of ethyl acetate to generate a turbid suspension. Then the 1-azido-3-propanol (2g, 15 mmol) was added into above turbid solution drop by drop until it became clear and further stirring for 4 h at room temperature. The rest of the liquid was extracted by ethyl acetate and washed by DI water for 3 times. The ethyl acetate solution containing product was dried using magnesium sulfate overnight as followed with rotary evaporator to get liquid product.

(Supplementary Fig. 3) $^1$H NMR (400 MHz, DMSO) δ 8.28 (dt, *J* = 15.1, 1.1 Hz, 1H), 7.60 (t, *J* = 1.4 Hz, 1H), 7.06 (d, *J* = 0.0 Hz, 1H), 4.45 (t, *J* = 6.1 Hz, 2H), 3.48 (dt, *J* = 50.2, 6.7 Hz, 2H), 2.04 – 1.97 (m, 2H). $^{13}$C NMR (101 MHz, DMSO) δ 148.64, 137.67, 130.63, 117.84, 65.74, 47.88, 27.81.

LC-MS (m/z): Calcd for $[M+H]^+$: 196.08; $[M+Na]^+$: 218.08. Found: 196.127, 218.125

***Synthesis of 2-(2-(2-azidoethoxy)ethoxy)ethyl-1H-imidazole-2-carboxylate (Azido-PEG-CDI).***

CDI (2.35 g, 14.5 mmol) was dissolved in 20 mL of ethyl acetate to generate a turbid suspension. Next, the 1-azido-3-propanol (1g, 3.7 mmol) was added into above turbid solution drop by drop until it became clear and further stirring for 4 h at room temperature. The rest of the liquid was extracted by ethyl acetate and washed by DI water for 3 times. The ethyl acetate solution containing product was dried using magnesium sulfate overnight as followed with rotary evaporator to get liquid product.

(Supplementary Fig. 4) $^1$H NMR (400 MHz, DMSO) δ 8.26 (t, *J* = 1.1 Hz, 1H), 7.60 (t, *J* = 1.5 Hz, 1H), 7.09 (dd, *J* = 1.7, 0.9 Hz, 1H), 4.54 – 4.49 (m, 2H), 3.80 – 3.77 (m, 2H), 3.62 – 3.57 (m, 6H), 3.36 (dd, *J* = 5.6, 4.2 Hz, 2H). $^{13}$C NMR (101 MHz, DMSO) δ 148.84, 137.67, 130.82, 117.96, 70.31, 70.12, 69.74, 68.35, 67.67, 50.43.

LC-MS (m/z): Calcd for $[M+H]^+$: 270.11; $[M+Na]^+$: 292.11. Found: 270.611, 292.611

***Synthesis of Azido-Modified Dextran.***

Dextran (2 g, 0.018 mmol) and azido-CDI or azido-PEG-CDI were dissolved in 20 mL of anhydrous dimethyl sulfoxide (DMSO) and stirred overnight at 50 °C in a nitrogen atmosphere. The reaction solution was subsequently transferred into a dialysis tube (Mw = 3.5 kDa) against DI for 3 days. Finally, the white fluffy product was obtained by lyophilizing.

***Synthesis of Acetalated Azide Dextran.***

Acetalation of azido linker modified Briefly, 0.7 g azide Dextran was reacted with pyridinium p-toluenesulfonate (15 mg, 0.08 mmol) and 2-methoxypropene (1.7 mL, 111 mmol) in 20 mL of anhydrous DMSO and stirred for 6 hours under nitrogen atmosphere. The reaction solution was precipitated with DI and centrifuged to obtain hydrophobic dextran. The white powder was obtained by Freeze-drying.

***Synthesis of spermine modified Dextran***

The spermine modified Dextran was obtained according to our previous work[25].

**Proteomics analysis**

NPs were resuspended in 20μL Lysis buffer (consisting of 6M Guanidinium Hydrochloride, 10 mM TCEP, 40 mM CAA, 50 mM HEPES pH8.5) and incubated at room temperature for 30 minutes. Samples were diluted in 40μL digestion buffer (10% Acetonitrile in 50mM HEPES pH 8.5) and incubated with 250ng LysC (MS grade, Wako) for 4 hours at 37°C, 750RPM. Samples were further diluted in 140μL digestion buffer and incubated for 18 h with 250 ng trypsin (MS grade, Sigma) at 37°C, 750RPM. After digestion, the samples were acidified by adding trifluoroacetic acid (TFA) to a final concentration of 1%. Samples were centrifuged at 10000 RCF and supernatants were transferred to clean Protein LoBind Eppendorf tubes. Supernatants were desalted on a SOLAμ SPE plate (HRP, Thermo)[26]. Between each application, the solvent was spun through by centuriation at 1500 RPM. For each sample, the filters were activated with 200ul of 100% Methanol, then 200ul of 80% Acetonitrile, 0.1% formic acid. The filters were subsequently equilibrated 2x with 200ul of 1% TFA, 3% Acetonitrile, after which the sample was loaded. After washing the tips twice with 200ul of 0.1% formic acid, the peptides were eluted into clean 0.5ml Eppendorf tubes using 40% Acetonitrile, 0.1% formic acid. The eluted peptides were concentrated in an Eppendorf Speedvac, and re-constituted in 12μL buffer A* (2% Acetonitrile, 1% TFA) containing iRT peptides (Biognosys). Peptide concentrations were determined by nanodrop (DeNovix). Secondly, peptides were loaded onto a 2cm C18 trap column (Thermo Fisher 164705), connected in-line to a 15 cm C18 reverse-phase analytical column (Thermo EasySpray ES803) using 100% Buffer A (0.1% Formic acid in water) at 750bar, using the Thermo EasyLC 1200 HPLC system, and the column oven operating at 35°C. Peptides were eluted over a 140 min gradient ranging from 6 to 60% of 80% acetonitrile, 0.1% formic acid at 250 nl/min, and the Q-Exactive instrument (Thermo Fisher Scientific) was run in a DD-MS2 top10 method. Full MS spectra were collected at a resolution of 70,000, with an AGC target of $3\times10^6$ or maximum injection time of 20 ms and a scan range of 300–1750 m/z. The MS2 spectra were obtained at a resolution of 17,500, with an AGC target value of $1\times10^6$ or maximum injection time of 60 ms, a normalised collision energy of 25 and an intensity threshold of $1.7e^4$. Dynamic exclusion was set to 60 s, and ions with a charge state <2 or unknown were excluded. MS performance was verified for consistency by running complex cell lysate quality control standards, and chromatography was monitored to check for reproducibility. Finally, all raw LC-MS/MS data files were processed together using Proteome Discoverer version 2.4 (Thermo) with the use of Label-free quantitation (LFQ) in both the processing and consensus steps. In the processing step, Oxidation (M), protein N-termini acetylation and met-loss were set as dynamic modifications, with cysteine carbamidomethyl set as static modification. All results were filtered with percolator using a 1% false discovery rate (FDR), and Minora Feature Detector was used for quantitation. SequestHT was used as for matching spectra against the human database from Uniprot.

### Supplementary figures

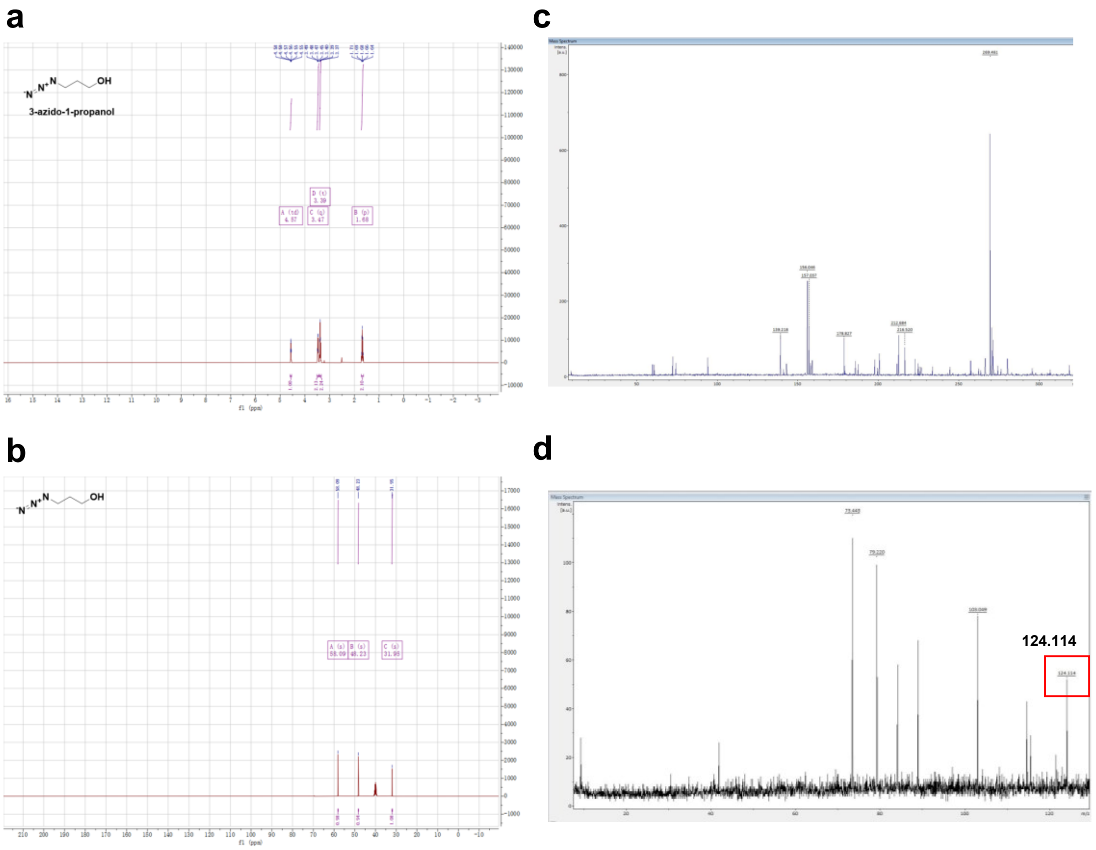

**Supplementary Fig. 1 | Nuclear magnetic resonance (NMR) Mass spectrometric quantification of 3-azido-1-propanol. (a)** $^{1}$HNMR, **(b)** $^{13}$CNMR of 3-azido-1-propanol, **(c)** MALDI-TOF mass spectrometry background of matrix DHB and **(d)** mass spectrum of 3-azido-1-propanol.

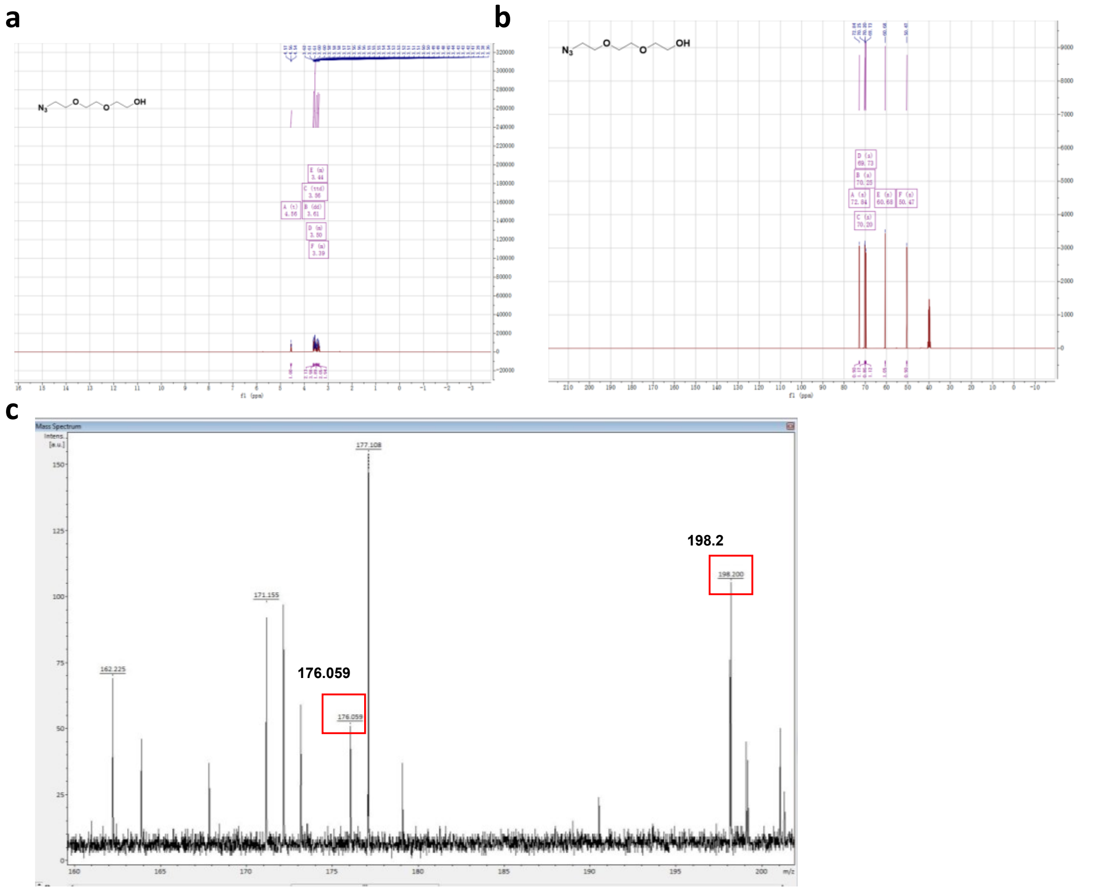

**Supplementary Fig. 2 | NMR and Mass spectrometric characterization of Azido-PEG3-OH. (a)** $^{1}$HNMR of Azido-PEG3-OH, **(b)**$^{13}$CNMR of Azido-PEG3-OH and **(c)** MALDI-TOF mass spectrometry of Azido-PEG3-OH.

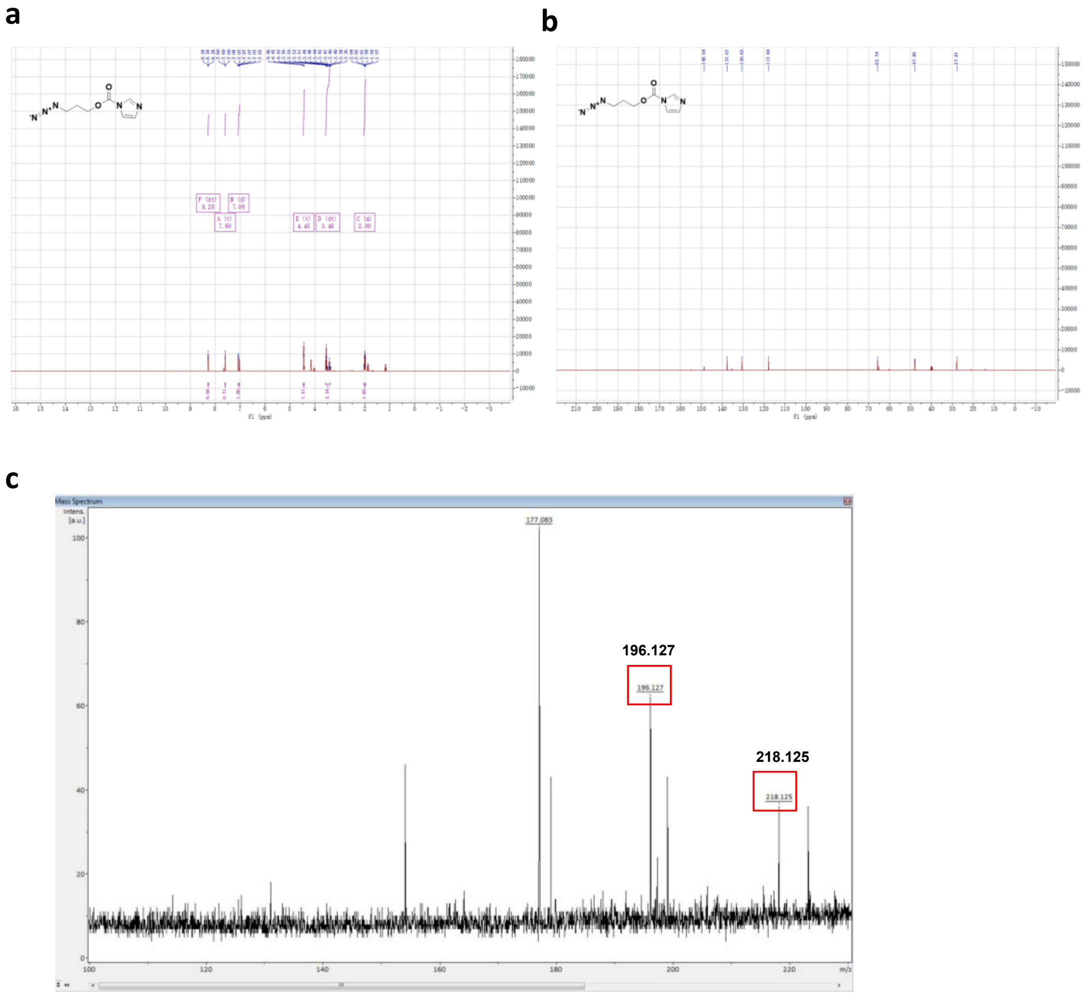

**Supplementary Fig. 3 | NMR and Mass spectrometric quantification of Azido-CDI.** **(a)** $^{1}$HNMR of Azido-CDI and **(b)**$^{13}$CNMR of Azido-CDI and **(c)** MALDI-TOF mass spectrometry of Azido-CDI.

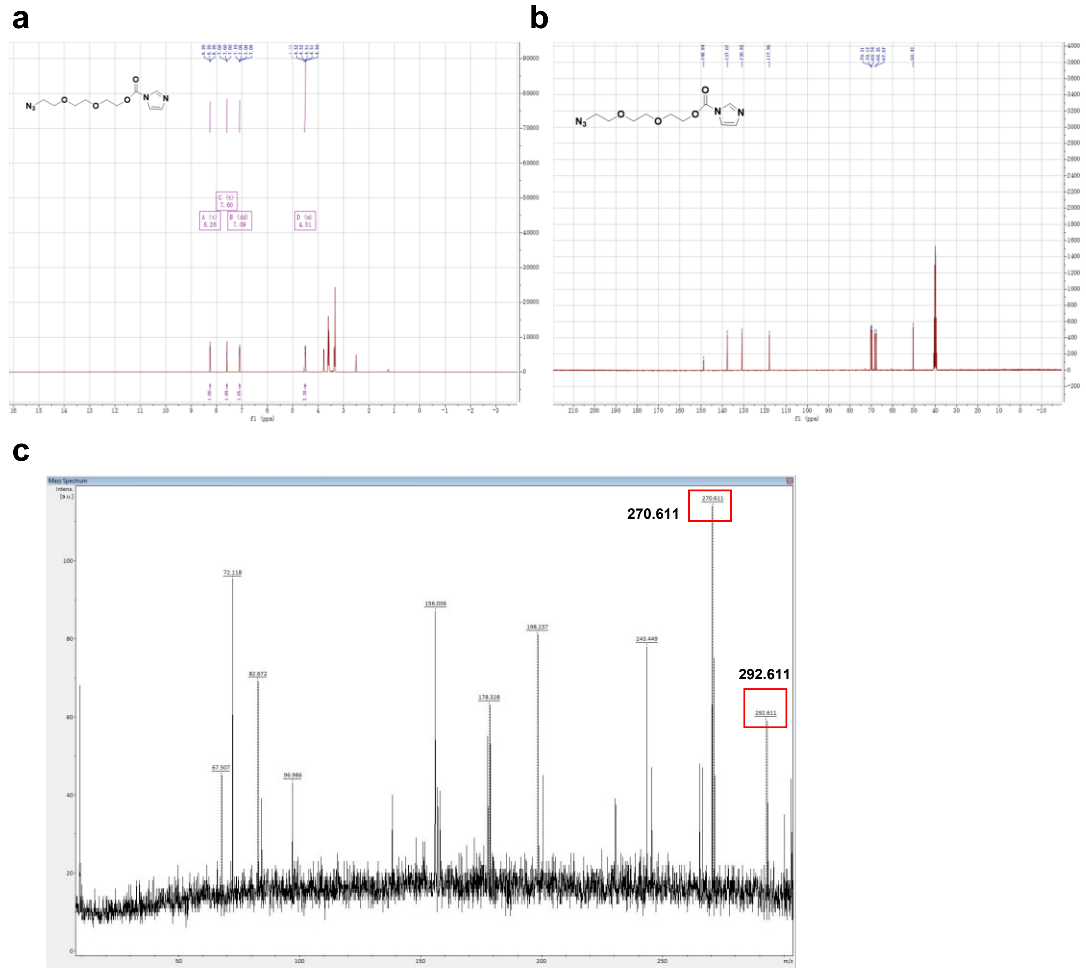

**Supplementary Fig. 4 | NMR and Mass spectrometric characterization of Azido-PEG3-CDI. (a)** $^{1}$HNMR of Azido-PEG3-CDI, (b) $^{13}$CNMR of AzidoPEG3-CDI and (c) MALDI-TOF mass spectrometry of Azido-PEG3-CDI.

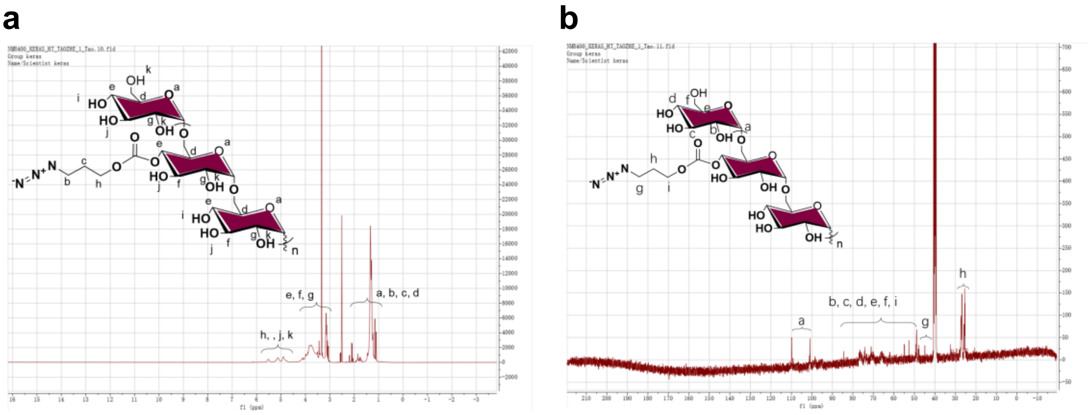

**Supplementary Fig. 5 | NMR characterization of azido-dextran. (a)** $^{1}$HNMR of azido-dextran. **(b)**$^{13}$CNMR of azido-dextran.

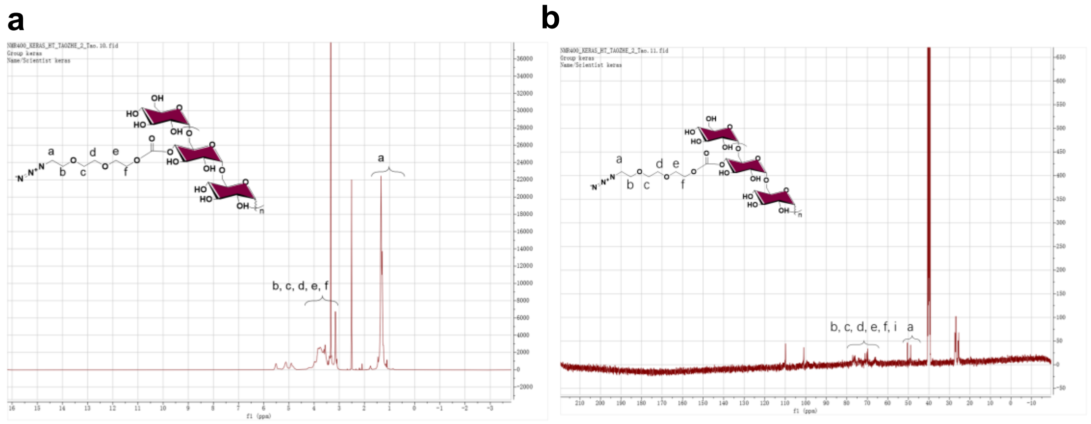

**Supplementary Fig. 6 | NMR characterization of azido-PEG3-dextran. (a)** $^{1}$HNMR of azido-PEG3-dextran. **(b)**$^{13}$CNMR of azido-PEG3-dextran.

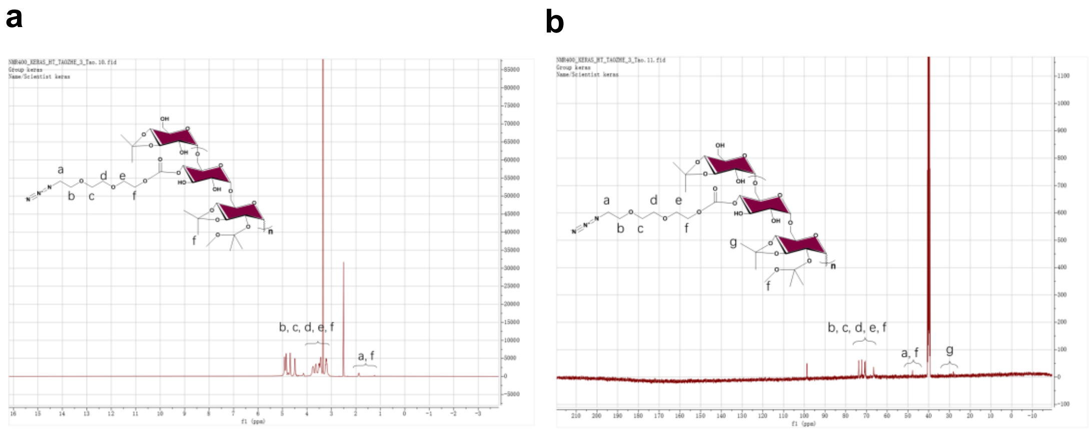

**Supplementary Fig. 7 | NMR characterization of acetalated azido-PEG3-dextran. (a)** $^{1}$HNMR of acetalated azido-PEG3-dextran. **(b)** $^{13}$CNMR of acetalated azido-PEG3-dextran.

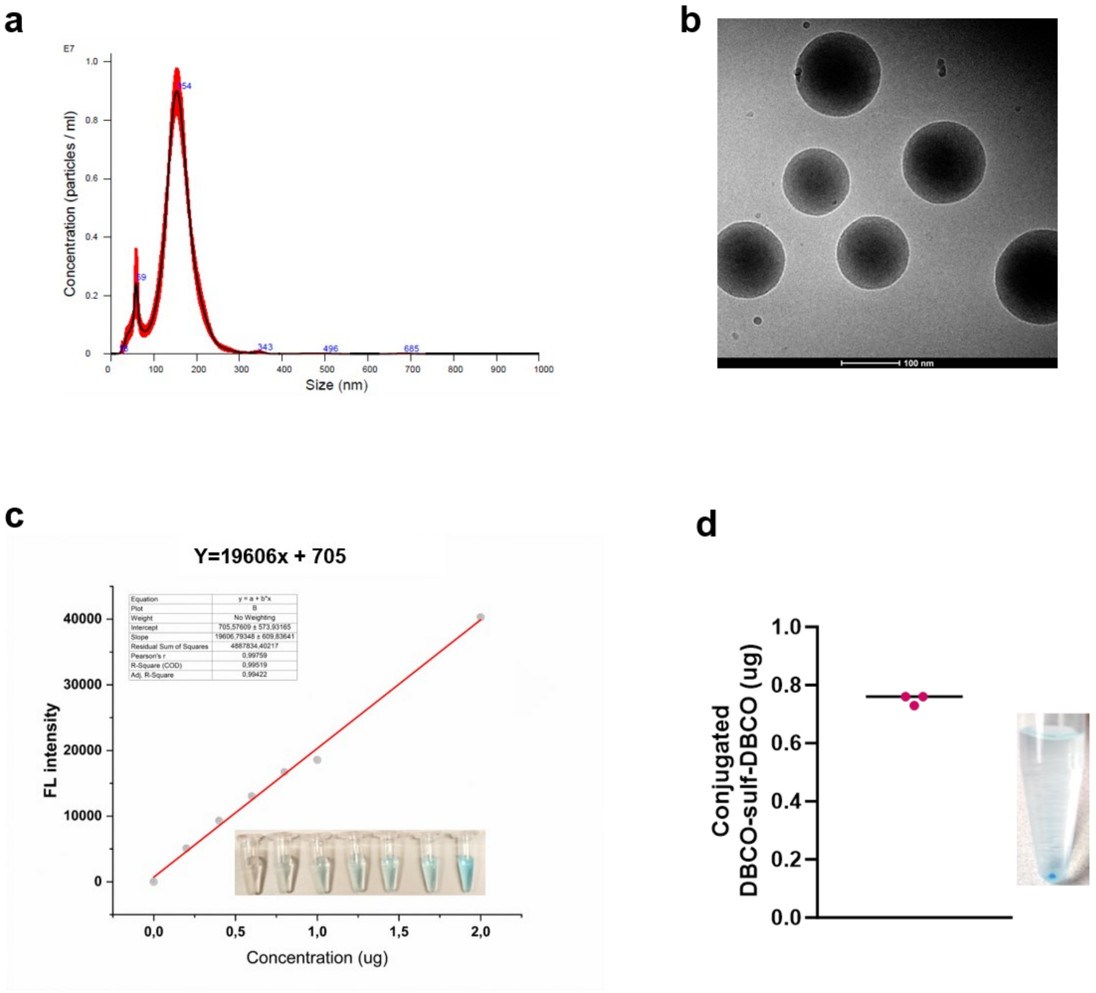

**Supplementary Fig. 8 | Characterization of dextran NPs (naked NPs) and the measurement of the concentration of available azido groups on surface of naked NPs. (a)** Size measurement of naked NPs by nanoparticle tracking analysis (n=5). **(b)** Cryo-TEM visualizing of naked NPs, scale bar: 100 nm. **(c)** Linear fitting curve of the concentration of dibenzocyclooctyne- Cyanine5 (DBCO-Cy5). **(d)**The density of azido groups available for clicks on the surface of dextran NPs.

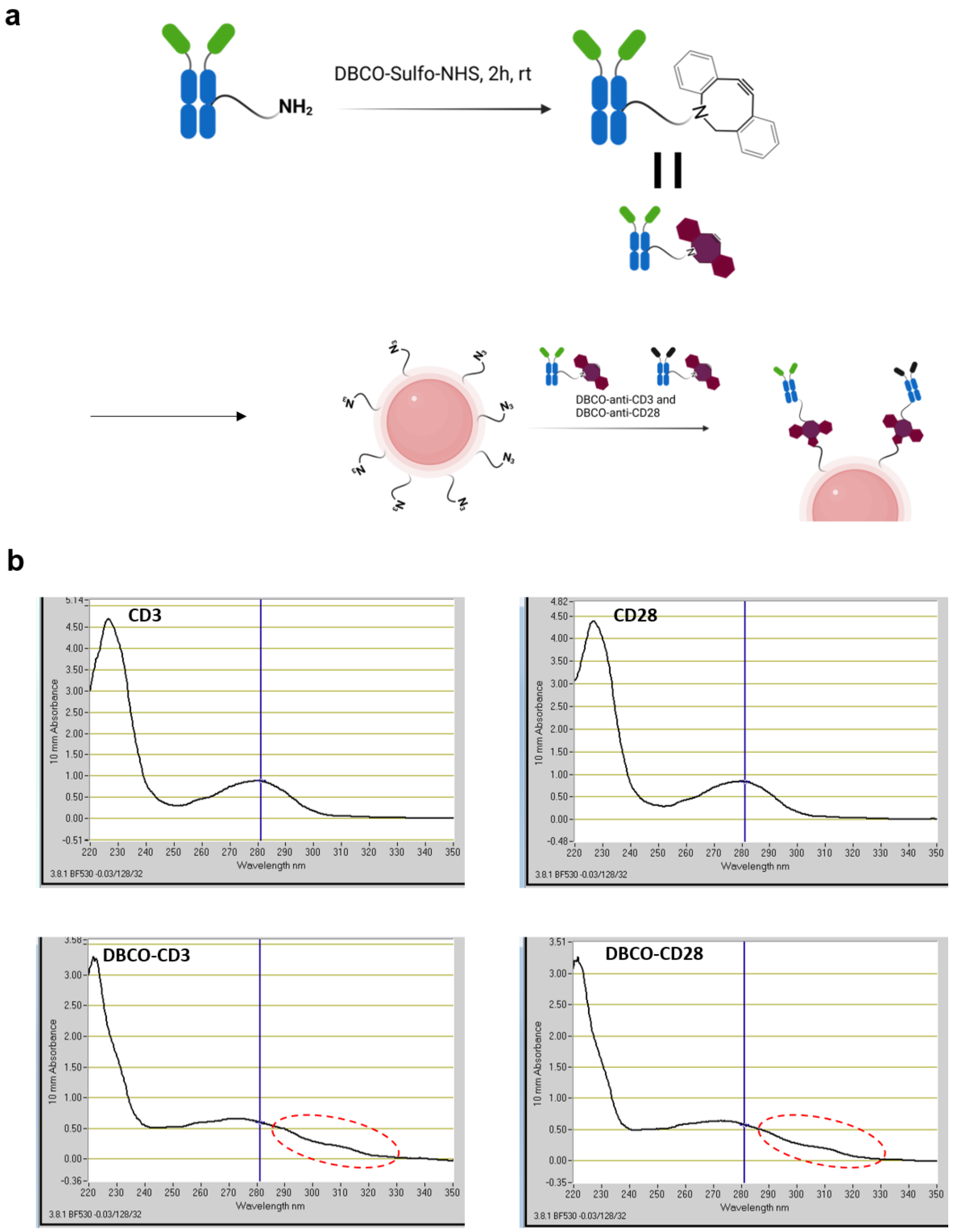

**Supplementary Fig. 9 | Illustration of antibody conjugated onto surface of naked NPs.** (a) schematic for conjugation of anti-CD3 and anti-CD28 antibodies with DBCO, then the DBCO modified antibody will direct conjugated onto the surface of naked NPs through click chemistry. **(b)** Nanodrop measurement of DBCO-conjugated ant-CD3 antibody and DBCO-conjugated anti-CD28 antibody showing distinct absorption peak of DBCO from 300-320 nm.

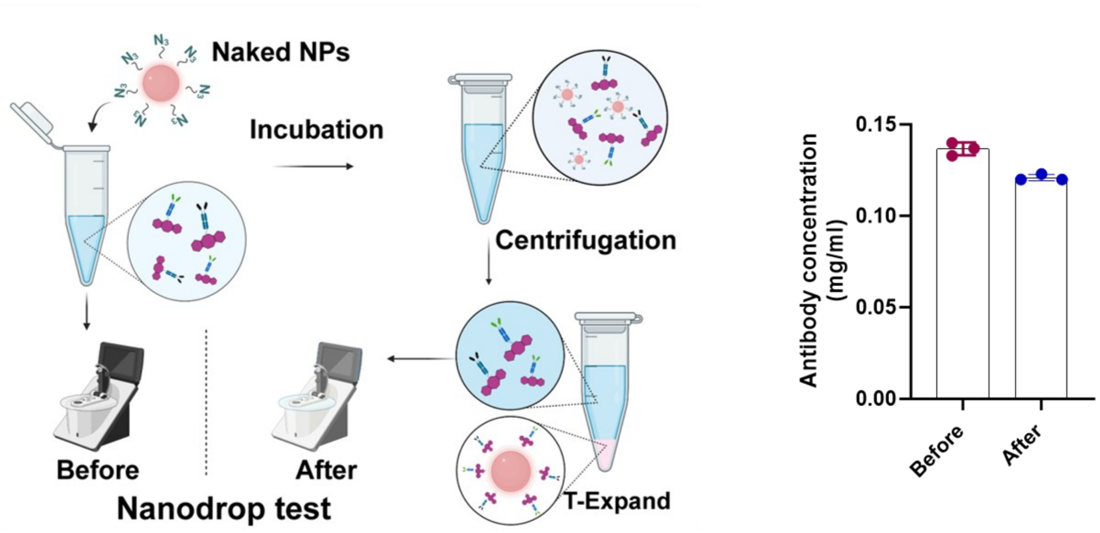

**Supplementary Fig. 10 | Illustration of antibody conjugation efficiency.** The antibody conjugation efficiency on the surface of naked NPs measured by Nanodrop.

**Proteomics analysis of T-Expand**

Click chemistry facilitates efficient antibody conjugation to NP surfaces, yet challenges persist when incubating the NPs with culturing medium. Protein corona formation in biological environments may obscure the antibodies, limiting their interactions with T cells. Hence, we conducted proteomics analysis to investigate protein corona formation on the T-Expand surface. Three types of the dextran NPs (A: amine-dextran NPs (positive charge surface); B: Azido-dextran NPs (negative charge surface, naked NPs); C: T-Expand) were incubated with full human serum, followed by centrifugation to remove any unbound protein (**Supplementary Fig. 11a**).

The composition of serum proteins absorbed on each particle were analyzed using shotgun proteomics [liquid chromatography tandem mass spectrometry (LC-MS/MS)]. A total of 666 proteins were identified across three groups (**Supplementary Fig. 11b**). The surface of cationic charge NPs (A) was enriched with a greater diversity of proteins with a total of 586 types detected, compared to 448 and 408 types found in B (naked NPs) and C (T-Expand), respectively **(Supplementary Fig. 11c)**. The overall protein profiles observed for NPs B and C were similar (**Supplementary Fig. 11d**), but the enriched proteins for T-Expand were less detected, indicating that the aCD3 and aCD28 conjugated on the naked NPs results in less interactions with serum proteins.

Among the top 20 abundant proteins which consisted of up to 90% of the total biomass (**Supplementary Table 2**). We found that T-Expand shows high level of serum albumin (**ALB, Supplementary Fig. 12a**), which will reduce the non-specific adsorption of NPs to other proteins, thereby enhancing the interaction between the T-Expand and T-cells to some extent[27]. Furthermore, we identified differential abundance proteins (DAPs) (**Supplementary Fig.12b, c**). Seven proteins showed significantly differential abundance across three groups (**Supplementary Fig. 12d**), in which C-reactive protein (**Supplementary Fig.12e**) and proprotein convertase subtilisin/kexin type 9 (**Supplementary Fig.12f**) showed significantly low abundance in group B and C compared with groups A. Meanwhile, we also observed a significantly high abundance of histidine-rich glycoprotein (**HRG**) in T-Expand (**Supplementary Fig.12g**), which likely will prevent T-Expand from being engulfed by macrophages in the body[28].

Next, we verified the ability of T-Expand to bind T-cells after forming the protein corona. We pre-incubated T-Expand with human serum and subsequently incubated with both Luciferase NFAT Jurkat cells, a cell line used to measure T-cell activation, and primary human T-cells (Supplementary Fig.10h). Notably, luciferase expression increased in correlation with T-Expand concentration, remaining unaffected by the presence of the protein corona (Supplementary Fig.10 i). Similarly, in the T-cell activation assay, we observed no attenuation in the activation signal (co-expression of CD25 and CD69 was 80%), indicating that both the functional orientation and antigen-binding capacity of the antibodies were preserved. Overall, the proteomic study results suggest that due to the unique material design and surface conjugation chemistry, the accessibility of T-Expand for T cell engagement is not affected by protein corona formed around the NPs.

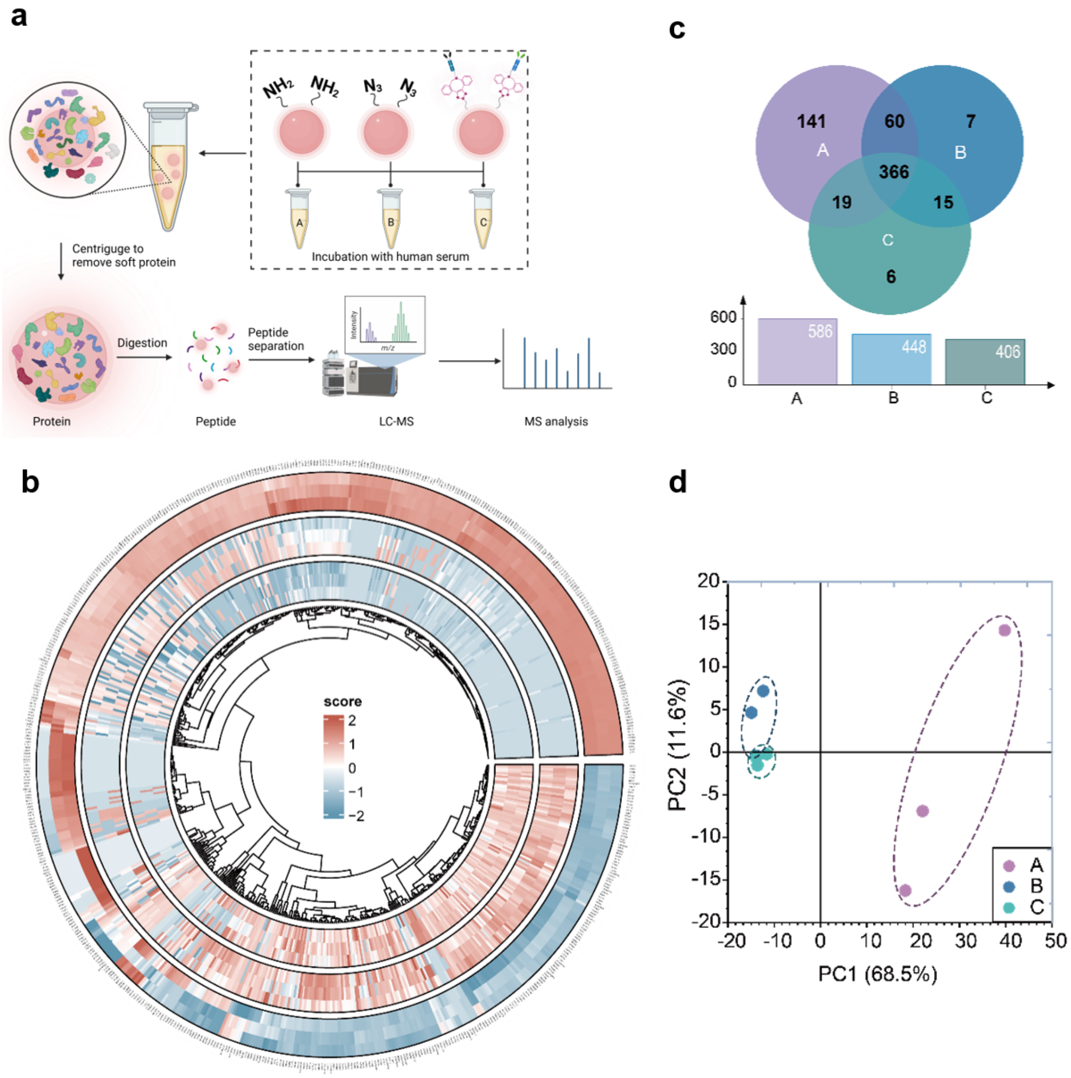

**Supplementary Fig. 11 | Proteomics analysis of protein corona formation on T-Expand. (a)** Illustration of the procedures of proteomics analysis to detect protein distribution on the surface of T-Expand. **(b)** Circular heatmap representing three different groups of the dataset after normalization and log2 transformation. Group A, B and C is visualized with a row-clustered heatmap, displayed on the outer, middle, and inner layers of the plot, respectively. The dendrogram was displayed by the protein abundance level of group C on the inside of the circular heatmap. The color gradient reflects normalized scores, with the legend indicating the corresponding values. **(c)** Protein corona components of three nanoparticles (venn plot & barplot) (A, B, C), **(d)** Principal Component Analysis (PCA) biplot of the dataset. The plot displays the first two principal components (PC1 and PC2), which account for the largest proportion of variance in the data. Each point represents a sample. The explained variance for PC1 and PC2 is provided in the axes labels.

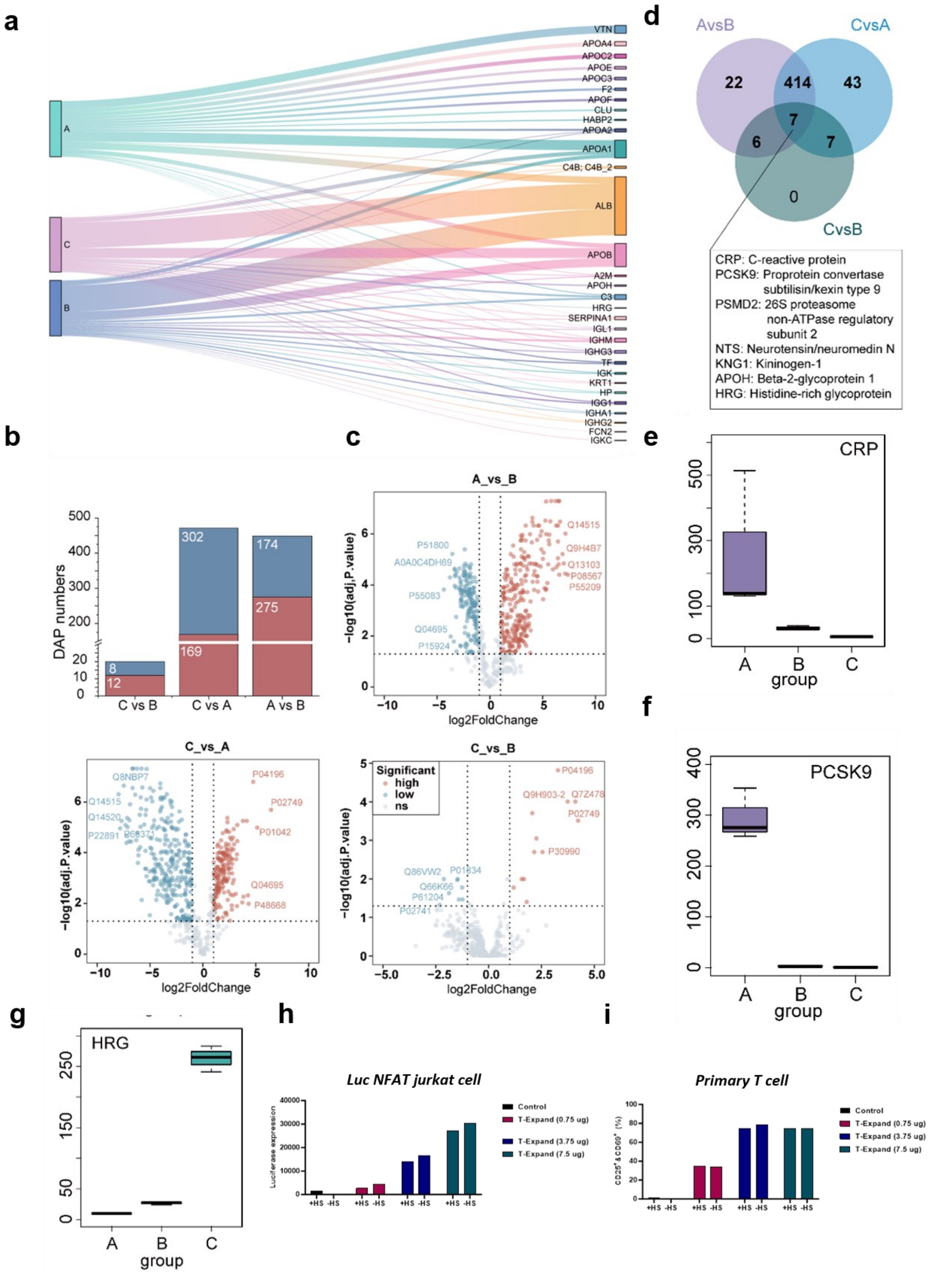

**Supplementary Fig. 12 | The accessibility of aCD3 and aCD28 on T-Expand for T-cell after incubation with human serum and formation of protein corona. (a)** The Sankey diagram of top 20 enriched protein among the three different groups of NPs. The width of each ribbon indicates the relative abundance represented in each group. (b) Numbers of the differential abundant proteins (DAPs) across three comparison groups. The high and low abundance were shown in red and blue, respectively. (c) Volcano plots of differentially abundant proteins in three comparisons. Proteins with significant differential abundance in each comparison are displayed in red and blue, with the top 5 upregulated (high) and top 5 downregulated (low) proteins annotated in the plot. (d) Common DAPs in the pairwise comparison across three groups (e-g) Boxplot showing the abundances levels of seven common differentially abundant s proteins across three comparisons: A vs. B, C vs. B, and C vs. A. Each box represents the distribution of scaled abundances after normalization for the respective proteins. The central line within each box indicates the median abundances level, while the box itself represents the interquartile range (IQR). Outliers are plotted as individual points. (h) Luciferase expression in NFAT Jurkat cells when incubated with different concentration of T-Expand pre-incubated with human serum for 24 h. (i) T cell activation marker (CD25 and CD69) expression determined by flow cytometry 24h after incubation with serum coated T-Expand.

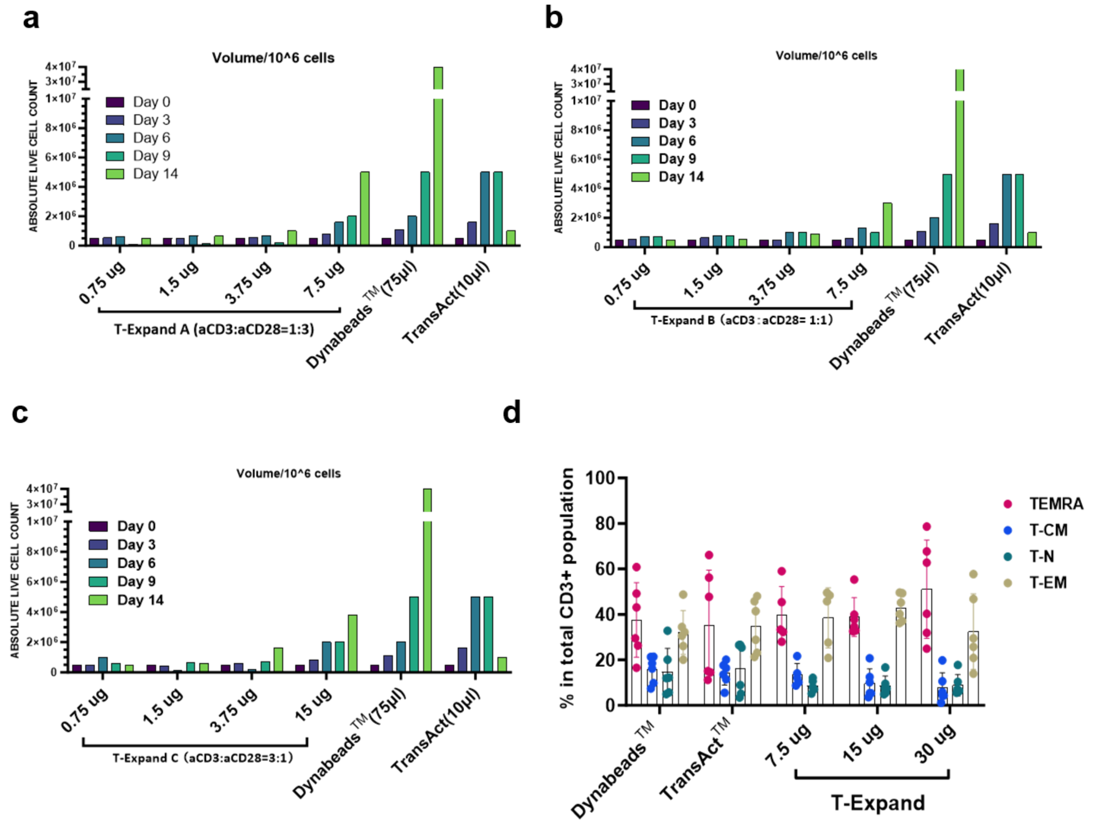

**Supplementary Fig. 13 | The different ratios (anti-CD3 : anti-CD28) on T-Expand on refers to T cell expansion at different dosage. (a)** T-Expand-A (1:3) **(b)** T-Expand-B (1:1), and **(c)** T-Expand-C (3:1), at various dose (0.75 ug, 1.5 ug, 3.75 ug and 15 ug per 10^6 Pan T cells) compared to Dynabeads$^{TM}$ or TransAct$^{TM}$. **(d)** Phenotypic distribution of T-cells based on CCR7 and CD45RA expression, showing TN (CD45RA$^{+}$CCR7$^{+}$), TCM (CD45RA$^{-}$CCR7$^{+}$), TEM (CD45RA-CCR7-), and TEMRA (CD45RA+CCR7-) subset, The T-Expand-expanded T-cells predominantly consisted of 35-50% effector memory T-cells (TEM) and approximately 40% effector memory T- cells re-expressing CD45RA (TEMRA), which mirrored the distribution observed in Dynabeads$^{TM}$-expanded T-cells.

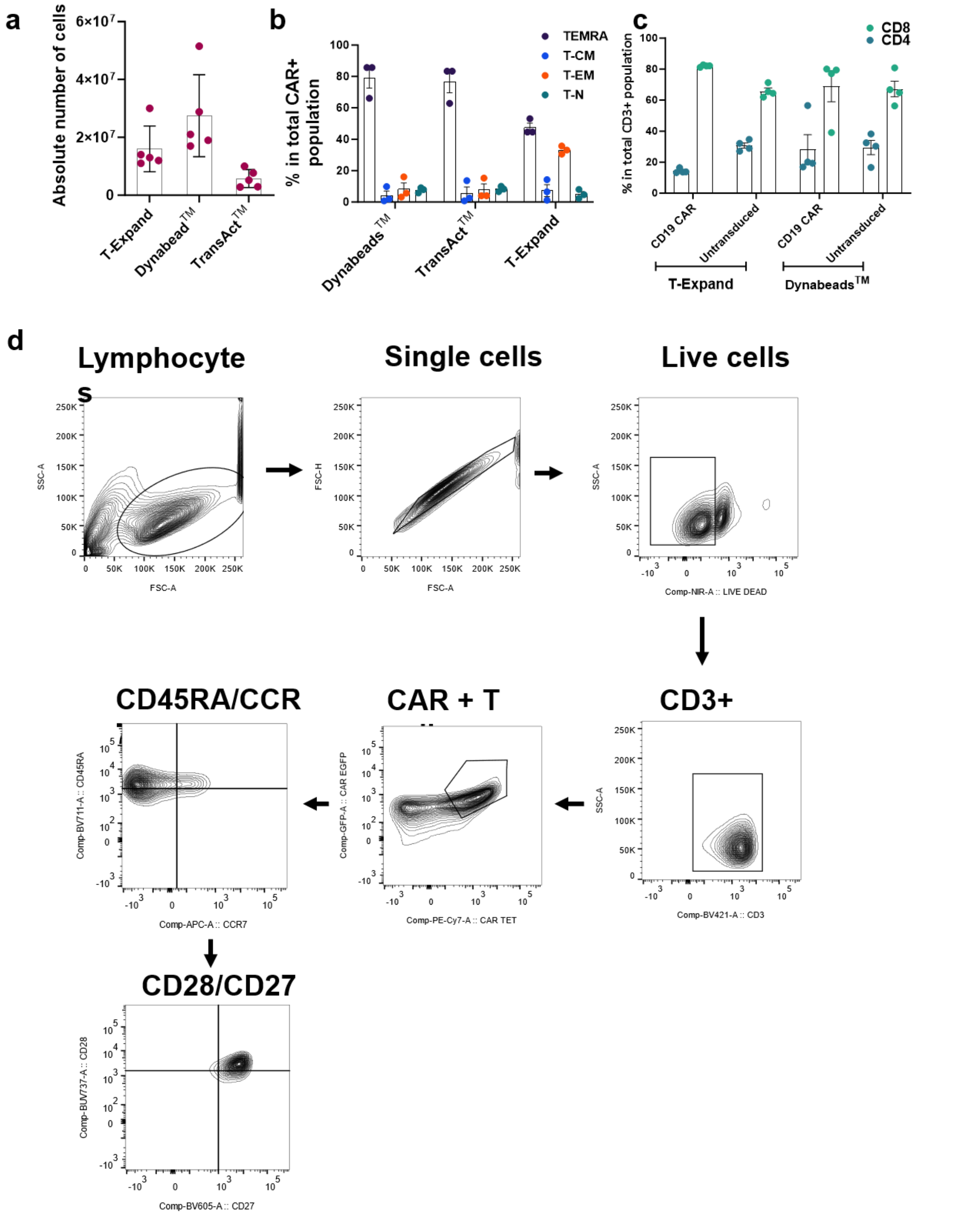

**Supplementary Fig. 14 | T-Expand mediated CD19 CAR T-expansion. (a)** Shows the absolute cell numbers of CAR T-cells over the expansion with T-Expand (or) Dynabeads™ (or) TransAct™. **(b)** Phenotypic distribution of T-cells based on CCR7 and CD45RA expression, showing TN (CD45RA$^+$CCR7$^+$), TCM (CD45RA$^-$CCR7$^+$), TEM (CD45RA-CCR7-), and TEMRA (CD45RA+CCR7-) subset total CD19 CAR T cells expanded with T-Expand (or) Dynabeads™ (or) TransAct™. **(c)** A graph showing the CD4+CD8+ T cells of total CD19 CAR T-cells expanded with T-Expand (or) Dynabeads™. **(d)** Gating strategy of CD19 CAR T-cell with representative flow plots of phenotype.

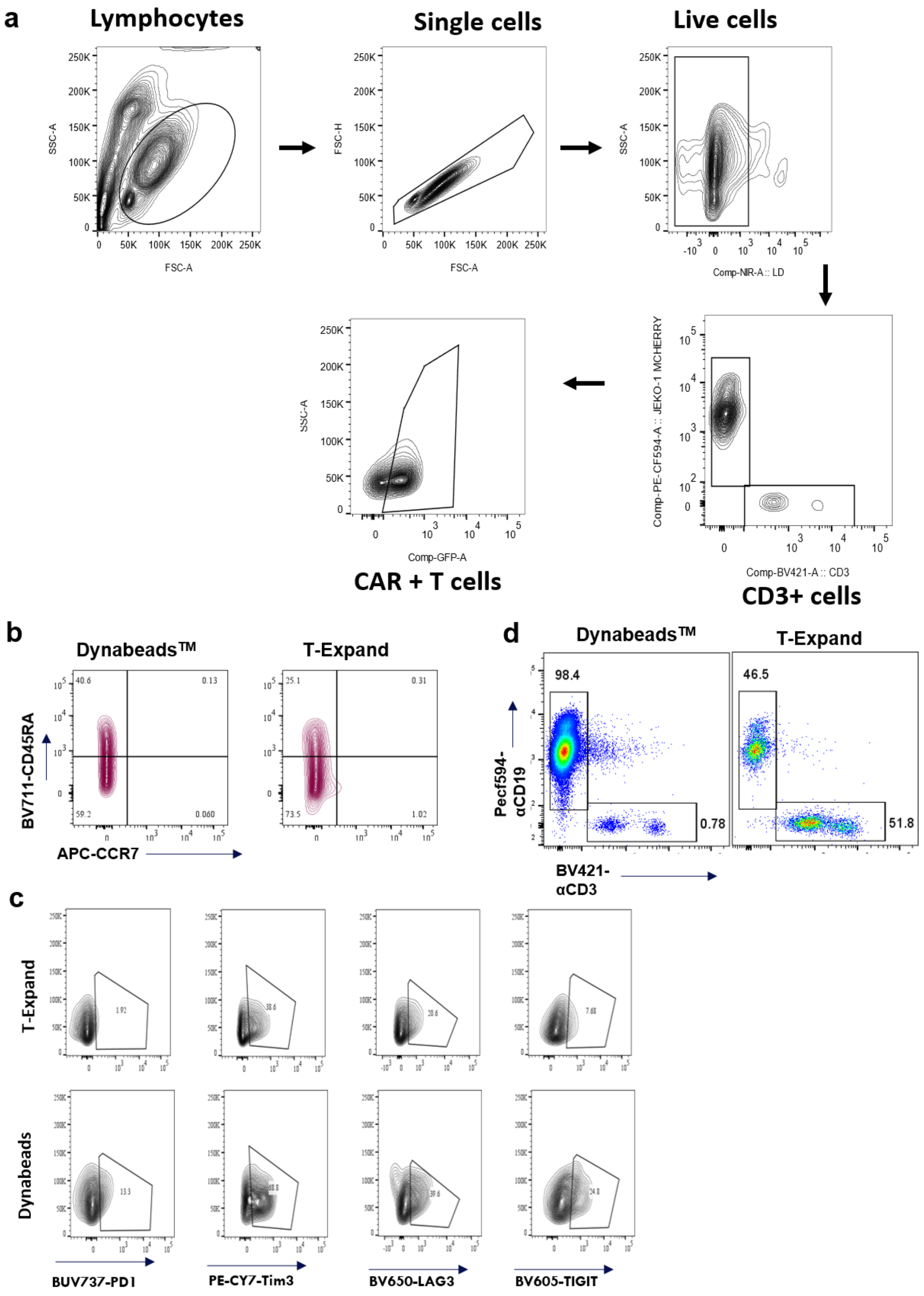

**Supplementary Fig. 15 | Gating strategy of rechallenge killing phenotype & representative flow plots of phenotype and exhaustion.** Flow cytometry gating strategy for CART cells after rechallenge. **(b)** Phenotypic distribution of CAR T-cells expanded with T-Expand (or) Dynabeads$^{TM}$ (or) TransAct$^{TM}$. based on CCR7 and CD45RA expression, showing TN (CD45RA$^{+}$CCR7$^{+}$), TCM (CD45RA$^{-}$CCR7$^{+}$), TEM (CD45RA-CCR7-), and TEMRA (CD45RA+CCR7-) subset **(c)** Flow cytometry analysis of the percentage of the CAR T cells expressing exhaustion markers such as PD1, LAG3, TIM3, and TIGIT. **(d)** Flow cytometry analysis showing the percentage of CD3+ T cells in the total live cell population at E/T ratio of 1:1 and 5:1 after three rounds of re-challenge.

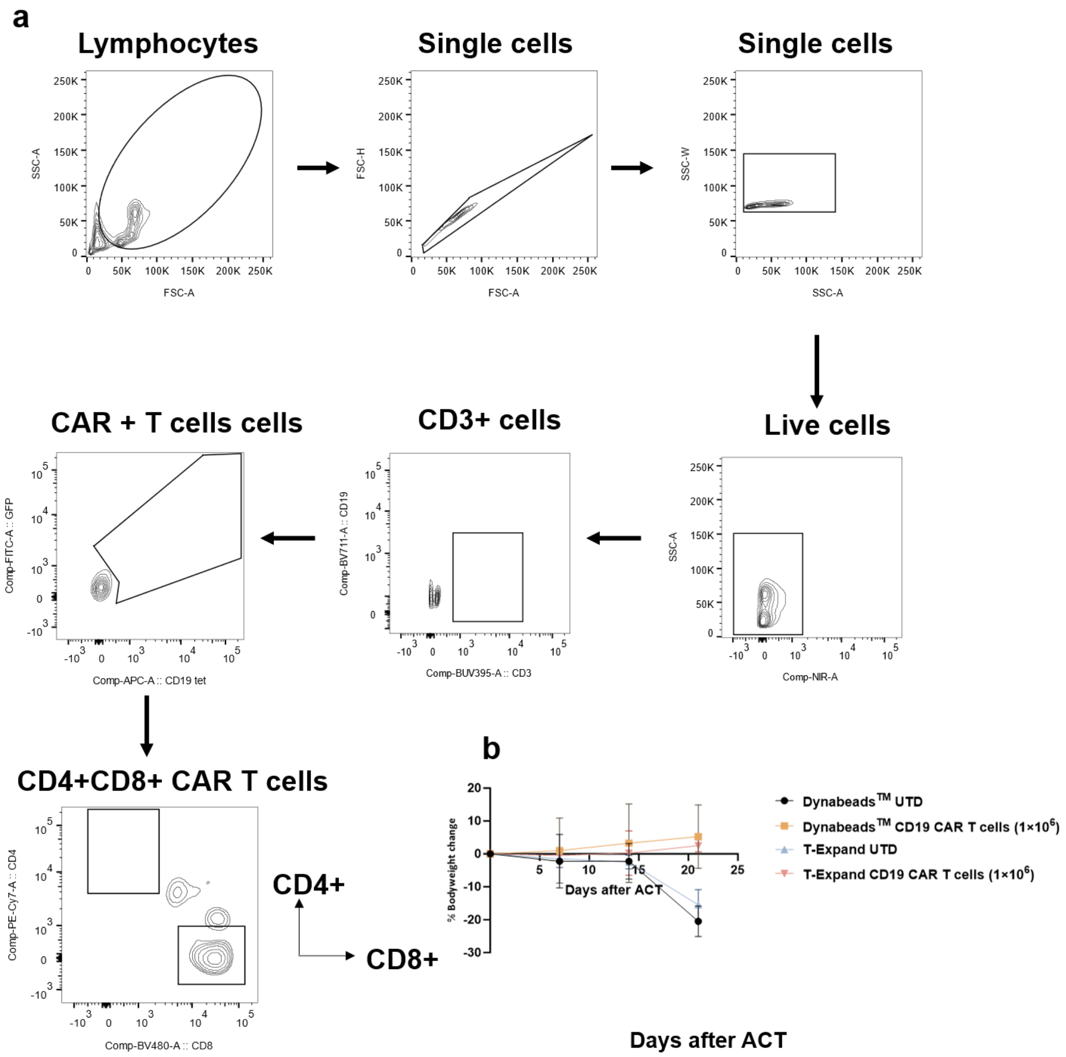

**Supplementary Fig. 16 | T-expand expanded CAR T display antitumor activity in a xenograft model of B-cell lymphoma.** (a) Gating strategy of mice spleen and bone marrow on Flowjo. (b) Percent change in body weight relative to pre-tumor injection in each group.

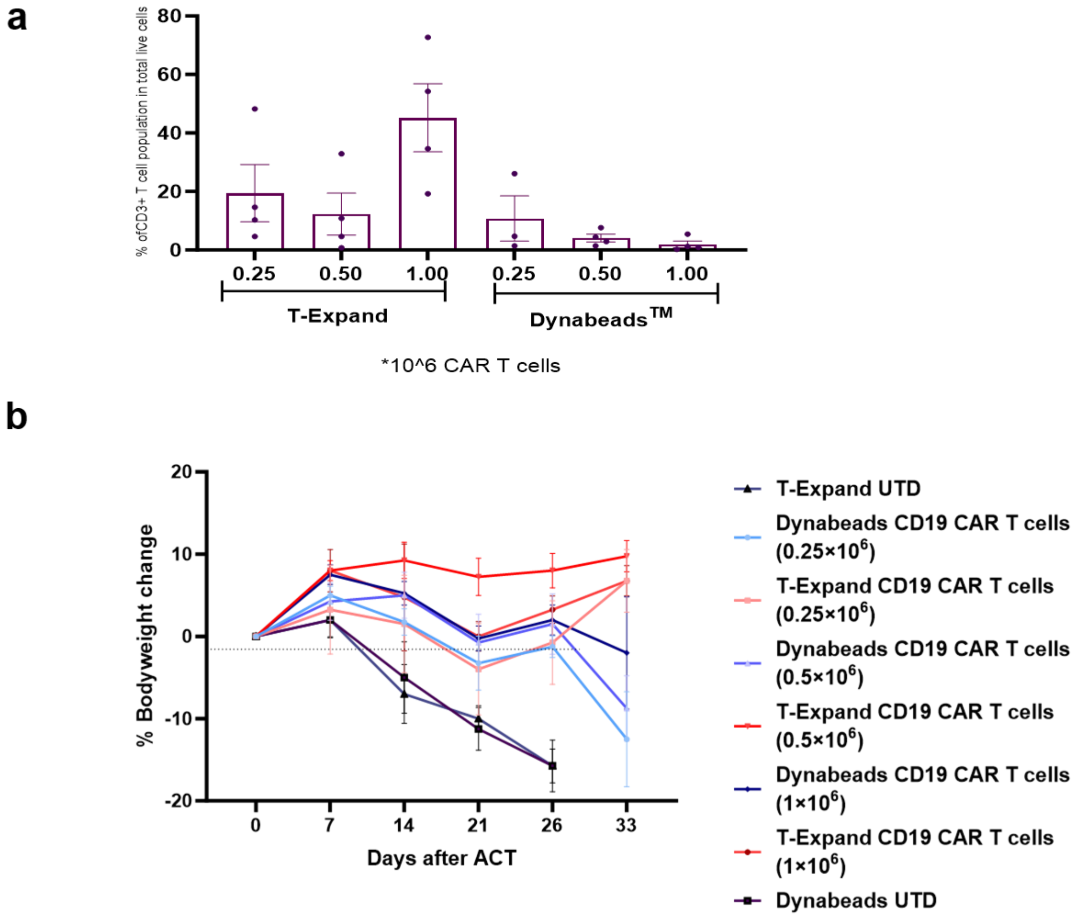

**Supplementary Fig. 17 | Dose de-escalation experiment to further elucidate differences between CAR T cells expanded with T-Expand or Dynabeads™.** (a) blood analysis of percentage of circulating CD3+ T-cells population in mice receiving CAR T-cells expanded with T-Expand at day treatment of 21. (b) Percent change in body weight relative to pre-tumor injection in each group.

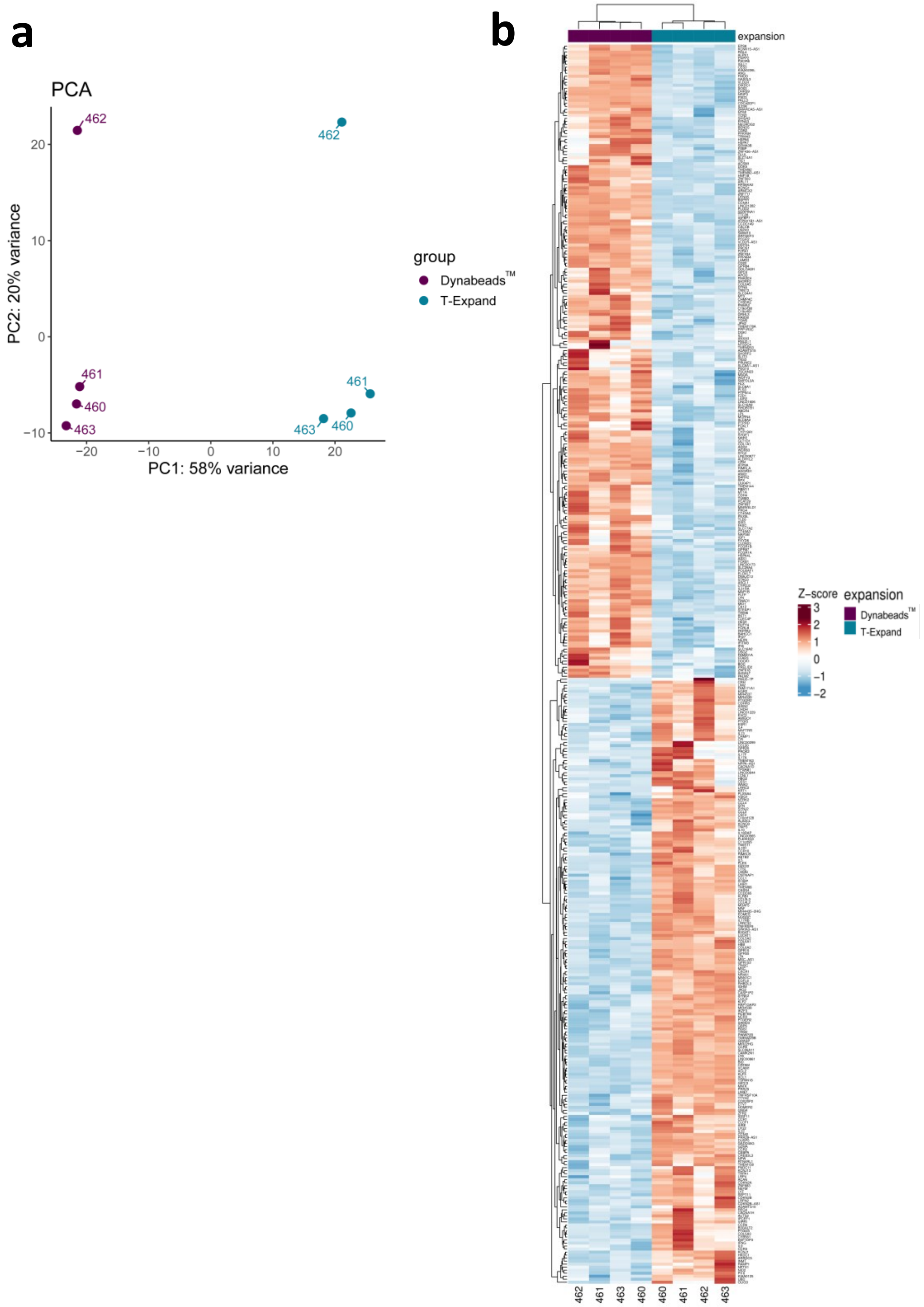

**Supplementary Fig. 18 | Deep RNA sequencing analysis of expanded CAR T cells expanded with T-Expand or Dynabeads™.** (a) The principal component analysis (PCA) revealed that the overall gene profiles of T-Expand expanded CAR T cells. (b) Unsupervised clustering of the most differentially expressed genes identified through differential gene expression analysis of CAR T cells expanded using either T-Expand or Dynabeads™.

### Supplementary tables

### Supplementary Table 1. Microscale- and Nanoscale-Based aAPCs for Polyclonal expansion of primary human or mouse T cells

| Microscaffolds | Size (μm) | Antibody conc | CAR T production | Ref |
|---|---|---|---|---|
| Dynabeads | 4.5 | Anti-CD3<br>Anti-CD28 | YES | 1, 2[1] |
| PLGA | 8 | Ratio of Anti-CD3 to Anti-CD28= 1:1<br>Conjugated concentration not mentioned | NO | 3 |
| Alginate microgel | 72 ± 2 nm | Ratio of Anti-CD3 to Anti-CD28= 1 to 7<br>Conjugated concentration is from 0.4 to 1.6μg/cm | NO | 4 |
| Lipid coated Silica microparticles | 8 | Ratio of Anti-CD3: Anti-CD28= 1:5<br>0.02-2 protein/lipid mol % | No | 5 |
| Lipid coated Silica microrods | 70 × 4.5 | Ratio of Anti-CD3: Anti-CD28= 1:1<br>0.01-1 protein/lipid mol % | YES | 6, 7, 8, 9, 10 |
| Polydopamine DNA microflowers | 1.83 ± 0.17 | Ratio of Anti-CD3: Anti-CD28= 1:1.80<br>0.91 ± 0.05 μg/107 particles and 1.64 ±<br>0.05 μg/1 × 107 particles | NO | 11 |
| Dendritic cell membrane coated PCL microfibril | 3.4 ± 1.6 μm in diameter and 55.5 ± 23.8 μm in length | Ratio of Anti-CD3: Anti-CD28= 1:2<br>0.0064nM stimuli/ug microfibril | NO | 12 |
| Gaphene oxide (GO) | 11.8 ± 2.6 | Ratio of Anti-CD3: Anti-CD28= 1:1 | YES | 13 |

| | | | | |
|---|---|---|---|---|
| | | ~1,119 molecules per µm2<br>GO | | |
| Alginate microsphere | 7-9 | Ratio of Anti-CD3:<br>Anti-CD28= 3:1<br>higher than $10^{5.9}$ molecules per microsphere | YES | 14 |
| polycaprolactone (PCL) 3D scaffold | 200 µm to 1000 um fiber spacing; Fiber diameter: ~12.8 µm ± 0.38 µm | 36 ± 0.88 molecules/µm2 | NO | 15 |

| **Nanoscaffolds** | **Size (nm)** | **Antibody conc** | **CAR T production** | **Ref** |
|---|---|---|---|---|
| TransAct™ | 100 | Not Mentioned | YES | 2,16,17 |
| Lipids | 189.7 ± 97.3 | Ratio of Anti-CD3:<br>Anti-CD28= 1:10<br>12.7 and 109 ng /uL of lipids solution. | YES | 18 |
| Streptavidin | 200 | Not Mentioned | YES | 19 |
| Poly(isocyano peptides) | Length: 150-200 | Anti-CD3 3-5 molecules per 150-200 nm of polymer | NO | 20 |
| Gellan gum | 150 | Ratio of Anti-CD3:<br>Anti-CD28= 1:2<br>(unit: 30µg/mg of Gellan gum) | NO | 3 |
| Gold Nanoparticles | 50 | Not Mentioned | NO | 21 |
| Poly(ethylene glycol)-block-poly(D,L-lactide),(PEG-PDLLA) | 160-2000 | Ratio of Anti-CD3:<br>Anti-CD28= 1:2<br>3.4 nm2/mL | NO | 22 |
| Coiled coil-protein | 705.9 ± 160.4 | Ratio of Anti-CD3:<br>Anti-CD28= 1:1<br>2.72 ± 0.81 ng/µg of ccNPs | NO | 23 |
| Single-Walled Carbon Nanotube | Not mentioned | Ratio of Anti-CD3:<br>Anti-CD28= 1:1<br>5µg Ab/mL of nanotube solution. | NO | 24 |
| PLGA | 130 | Anti-CD3<br>Anti-CD28<br>1µg/mg polymer | NO | 3 |

### Supplementary Table 2. The twenty most abundant proteins in A, B and C samples

| | Group A | Protein | Percentage |
|---|---|---|---|
| 1 | P02647 | Apolipoprotein A-I OS=Homo sapiens GN=APOA1 PE=1 SV=1 | 16.02% |
| 2 | P04004 | Vitronectin OS=Homo sapiens GN=VTN PE=1 SV=1 | 12.51% |
| 3 | P02768 | Serum albumin OS=Homo sapiens GN=ALB PE=1 SV=2 | 10.80% |
| 4 | P04114 | Apolipoprotein B-100 OS=Homo sapiens GN=APOB PE=1 SV=2 | 7.02% |
| 5 | P06727 | Apolipoprotein A-IV OS=Homo sapiens GN=APOA4 PE=1 SV=3 | 6.71% |
| 6 | P02655 | Apolipoprotein C-II OS=Homo sapiens GN=APOC2 PE=1 SV=1 | 6.44% |
| 7 | P02649 | Apolipoprotein E OS=Homo sapiens GN=APOE PE=1 SV=1 | 4.16% |
| 8 | P02656 | Apolipoprotein C-III OS=Homo sapiens GN=APOC3 PE=1 SV=1 | 3.93% |
| 9 | P00734 | Prothrombin OS=Homo sapiens GN=F2 PE=1 SV=2 | 3.67% |
| 10 | Q13790 | Apolipoprotein F OS=Homo sapiens GN=APOF PE=1 SV=2 | 3.62% |
| 11 | P02652 | Apolipoprotein A-II OS=Homo sapiens GN=APOA2 PE=1 SV=1 | 3.20% |
| 12 | P10909 | Clusterin OS=Homo sapiens GN=CLU PE=1 SV=1 | 2.71% |
| 13 | Q14520 | Hyaluronan-binding protein 2 OS=Homo sapiens GN=HABP2 PE=1 SV=1 | 1.86% |
| 14 | P01024 | Complement C3 OS=Homo sapiens GN=C3 PE=1 SV=2 | 1.57% |
| 15 | P0C0L5 | Complement C4-B OS=Homo sapiens GN=C4B PE=1 SV=2 | 1.34% |
| 16 | P01871 | Immunoglobulin heavy constant mu OS=Homo sapiens GN=IGHM PE=1 SV=4 | 0.93% |
| 17 | P01009 | Alpha-1-antitrypsin OS=Homo sapiens GN=SERPINA1 PE=1 SV=3 | 0.78% |
| 18 | P01023 | Alpha-2-macroglobulin OS=Homo sapiens GN=A2M PE=1 SV=3 | 0.74% |
| 19 | P0DOX8 | Immunoglobulin lambda-1 light chain OS=Homo sapiens PE=1 SV=1 | 0.57% |
| 20 | P01860 | Immunoglobulin heavy constant gamma 3 OS=Homo sapiens GN=IGHG3 PE=1 SV=2 | 0.55% |
| | Total | | 89.14% |

| | Group B | Protein | Percentage |
|---|---|---|---|
| 1 | P02768 | Serum albumin OS=Homo sapiens GN=ALB PE=1 SV=2 | 42.01% |
| 2 | P04114 | Apolipoprotein B-100 OS=Homo sapiens GN=APOB PE=1 SV=2 | 13.95% |
| 3 | P02647 | Apolipoprotein A-I OS=Homo sapiens GN=APOA1 PE=1 SV=1 | 5.86% |
| 4 | P01871 | Immunoglobulin heavy constant mu OS=Homo sapiens GN=IGHM PE=1 SV=4 | 3.21% |
| 5 | P01024 | Complement C3 OS=Homo sapiens GN=C3 PE=1 SV=2 | 3.07% |
| 6 | P0DOX5 | Immunoglobulin gamma-1 heavy chain OS=Homo sapiens PE=1 SV=1 | 2.27% |
| 7 | P01009 | Alpha-1-antitrypsin OS=Homo sapiens GN=SERPINA1 PE=1 SV=3 | 2.24% |
| 8 | P02787 | Serotransferrin OS=Homo sapiens GN=TF PE=1 SV=3 | 2.12% |
| 9 | P01860 | Immunoglobulin heavy constant gamma 3 OS=Homo sapiens GN=IGHG3 PE=1 SV=2 | 2.09% |
| 10 | P0DOX8 | Immunoglobulin lambda-1 light chain OS=Homo sapiens PE=1 SV=1 | 1.56% |
| 11 | P00738 | Haptoglobin OS=Homo sapiens GN=HP PE=1 SV=1 | 1.48% |
| 12 | P0DOX7 | Immunoglobulin kappa light chain OS=Homo sapiens PE=1 SV=1 | 1.42% |
| 13 | P0C0L5 | Complement C4-B OS=Homo sapiens GN=C4B PE=1 SV=2 | 1.22% |
| 14 | P01876 | Immunoglobulin heavy constant alpha 1 OS=Homo sapiens GN=IGHA1 PE=1 SV=2 | 1.21% |
| 15 | P01023 | Alpha-2-macroglobulin OS=Homo sapiens GN=A2M PE=1 SV=3 | 1.06% |
| 16 | P01859 | Immunoglobulin heavy constant gamma 2 OS=Homo sapiens GN=IGHG2 PE=1 SV=2 | 0.96% |
| 17 | P02652 | Apolipoprotein A-II OS=Homo sapiens GN=APOA2 PE=1 SV=1 | 0.95% |
| 18 | P04264 | Keratin, type II cytoskeletal 1 OS=Homo sapiens GN=KRT1 PE=1 SV=6 | 0.83% |
| 19 | Q15485 | Ficolin-2 OS=Homo sapiens GN=FCN2 PE=1 SV=2 | 0.74% |
| 20 | P01834 | Immunoglobulin kappa constant OS=Homo sapiens GN=IGKC PE=1 SV=2 | 0.51% |
| | Total | | 88.74% |

| | Group C | Protein | Percentage |
|---|---|---|---|
| 1 | P02768 | Serum albumin OS=Homo sapiens GN=ALB PE=1 SV=2 | 40.52% |
| 2 | P04114 | Apolipoprotein B-100 OS=Homo sapiens GN=APOB PE=1 SV=2 | 16.06% |
| 3 | P02647 | Apolipoprotein A-I OS=Homo sapiens GN=APOA1 PE=1 SV=1 | 6.40% |
| 4 | P01024 | Complement C3 OS=Homo sapiens GN=C3 PE=1 SV=2 | 3.18% |
| 5 | P02749 | Beta-2-glycoprotein 1 OS=Homo sapiens GN=APOH PE=1 SV=3 | 2.20% |
| 6 | P01871 | Immunoglobulin heavy constant mu OS=Homo sapiens GN=IGHM PE=1 SV=4 | 2.13% |
| 7 | P02787 | Serotransferrin OS=Homo sapiens GN=TF PE=1 SV=3 | 2.09% |
| 8 | P01009 | Alpha-1-antitrypsin OS=Homo sapiens GN=SERPINA1 PE=1 SV=3 | 2.03% |
| 9 | P0DOX5 | Immunoglobulin gamma-1 heavy chain OS=Homo sapiens PE=1 SV=1 | 1.74% |
| 10 | P01860 | Immunoglobulin heavy constant gamma 3 OS=Homo sapiens GN=IGHG3 PE=1 SV=2 | 1.70% |
| 11 | P0DOX7 | Immunoglobulin kappa light chain OS=Homo sapiens PE=1 SV=1 | 1.33% |
| 12 | P00738 | Haptoglobin OS=Homo sapiens GN=HP PE=1 SV=1 | 1.23% |
| 13 | P0DOX8 | Immunoglobulin lambda-1 light chain OS=Homo sapiens PE=1 SV=1 | 1.06% |
| 14 | P0C0L5 | Complement C4-B OS=Homo sapiens GN=C4B PE=1 SV=2 | 1.02% |
| 15 | P01876 | Immunoglobulin heavy constant alpha 1 OS=Homo sapiens GN=IGHA1 PE=1 SV=2 | 0.94% |
| 16 | P02652 | Apolipoprotein A-II OS=Homo sapiens GN=APOA2 PE=1 SV=1 | 0.90% |
| 17 | P01023 | Alpha-2-macroglobulin OS=Homo sapiens GN=A2M PE=1 SV=3 | 0.79% |
| 18 | P04264 | Keratin, type II cytoskeletal 1 OS=Homo sapiens GN=KRT1 PE=1 SV=6 | 0.72% |
| 19 | P04196 | Histidine-rich glycoprotein OS=Homo sapiens GN=HRG PE=1 SV=1 | 0.66% |
| 20 | P01859 | Immunoglobulin heavy constant gamma 2 OS=Homo sapiens GN=IGHG2 PE=1 SV=2 | 0.65% |
| | Total | | 87.36% |

### Supplementary Table 3: antibody and cytokine information

| Antibodies& Protein | Manufacturer | Clone (Cat number) |
|---|---|---|
| anti-human CD3 | BioXcell | OKT-3 (BE0001-2) |
| anti-human CD28 | BioXcell | 9.3 (BE0248) |
| BB700-anti-human CD25 (IL-2R) | BD Biosciences | M-A251 (566447) |
| BV786-anti-human CD69 | BD Biosciences | FN50 (563834) |
| PE-anti-human CD69 | Biolegend | FN50 (310906) |
| BV421-anti-human CD3 | BD Biosciences | SP34-1 (562877) |
| BV711-anti-human CD45RA | BD Biosciences | HI100 (563733) |
| APC-anti-human CD197 (CCR7) | BD Biosciences | 2-L1-A (566762) |
| PE/Cy7-anti-human TIM-3 | Biolegend | F38-2E2 (345014) |
| BUV737-anti-human 279 (PD-1) | BD Biosciences | EH12.1 (612791, 565299) |
| BV650-anti-human CD223 (LAG-3) | BD Biosciences | 11C3C65 (369316) |
| PE-Cy7-anti-human CD57 | Biolegend | QA17A04 (393310, B323195) |
| BV786-anti-human CD39 | BD Biosciences | TU66 (742523) |
| BV605-anti-human TIGHT | BD Biosciences | 741182 (RUO) (747841) |
| BV480-anti-human CD4 | BD Biosciences | L200 (566148) |
| PerCP-anti-human CD8 | BD Biosciences | SK1 (347314) |
| BV480-anti-human CD8 | BD Biosciences | RPA-T8 (566121) |
| BUV737-anti-human CD28 | BD Biosciences | CD28.2 (612815) |
| BV605-anti-human CD27 | Biolegend | O323 (302830) |
| PE-anti-human CD137 | BD Biosciences | C65-485 (RUO) |
| PE/Cy7-Streptavidin | BD Biosciences | Cat: 557598 |
| BV711-anti-CD19 | BD Biosciences | SJ25C1 (563036) |
| PE-Cy7-anti-CD4 | BD Biosciences | RPA-T4 (560649) |
| Biotinylated Human CD19 (20-291) Protein | Acro Biosystems | 20-291(His,Avitag™, premium grade) |
| Human IL-2 Recombinant Protein, PeproTech® | Thermo Fisher Scientific | 200-02-50UG |

### Captions for the supplementary videos

| Video | Caption |
|---|---|
| Supplementary Video 1 | Airyscan imaging (3D view) on how T-Expand specifically interacting in engineered Jurkats, |
| Supplementary Video 2 | Airyscan imaging (3D view) on how Dynabeads™ interacting engineered Jurkats, |
